# Tuning the stator subunit of the flagellar motor with coiled-coil engineering

**DOI:** 10.1101/2023.03.05.530362

**Authors:** Pietro Ridone, Daniel L. Winter, Matthew A. B. Baker

## Abstract

Many bacteria swim driven by an extracellular filament rotated by the bacterial flagellar motor. This motor is powered by the stator complex, MotA_5_MotB_2_, a heterodimeric complex which forms an ion channel which couples energy from the ion motive force to torque generation. Recent structural work revealed that stator complex consists of a ring of five MotA subunits which rotate around a central dimer of MotB subunits. Transmembrane (TM) domains TM3 and TM4 from MotA combine with the single TM domain from MotB to form two separate ion channels within this complex. Much is known about the ion binding site and ion specificity; however, to date, no modelling has been undertaken to explore the MotB-MotB dimer stability and the role of MotB conformational dynamics during rotation. Here, we modelled the central MotB dimer using coiled-coil engineering and modelling principles and calculated free energies to identify stable states in the operating cycle of the stator. We found 3 stable coiled-coil states with dimer interface angles of 28°, 56° and 64°. We tested the effect of strategic mutagenesis on the comparative energy of the states and correlated motility with a specific hierarchy of stability between the three states. In general, our results indicate agreement with existing models describing a 36° rotation step of the MotA pentameric ring during the power stroke and provide an energetic basis for the coordinated rotation of the central MotB dimer based on coiled-coil modelling.

## Introduction

The bacterial flagellar motor (BFM) is a complex nanoscale engine that converts ion flux into the rotation of a flagellum to propel bacteria through their surroundings^1,2^. A crucial component of this complex is the stator subunit which is responsible for the torque-generating step of the motor^3,4^. In *E. coli*, the stator subunit is in fact composed of two transmembrane (TM) proteins (MotA and MotB) which assemble around the basal rotor of the flagellar motor and form a proton channel at their interface^5^. The inward flux of ions across the complex is efficiently coupled to a mechanical stimulus applied onto the rotor protein FliG which ultimately drives the rotation of a single flagellum^6^.

The structure of the full MotA_5_MotB_2_ complex was solved in 2020^7,8^, revealing that the stator complex likely acted as a rotary motor itself. Further recent structural work has identified the basis for ion selectivity and many catalytic residues involved in the process^9^. The current consensus model is that a transmembrane pentameric ring of MotA subunits rotates around a central dimer stalk of MotB TM domains^9,10^. The model details the conformational waypoints through which the MotA subunit passes during the gating cycle of the stator, but our understanding of the dynamics, in general, is limited, and in particularly for the MotB subunit central stalk.

The central MotB dimer is important because it harbours the catalytic centre. It has been proposed to rotate in place between protonated and non-protonated states^10^. Previous work suggested that the MotB TM domains exchange their interface contacts during gating and may pivot along their helical axis^11^, in a similar fashion to the proposed mechanism for the homologous TolR protein^12^. The dimerization of MotB at the TM domain was shown to be essential for motility as modifications to TM residues that restrict or alter the conformational space of the dimer (e.g., cysteine crosslinking and tryptophan-scanning mutagenesis) generally diminished motility^11,12^.

The MotB stalk appears to assemble as a dimeric parallel coiled-coil in recently published structures^7,8^. Coiled-coils are well-characterized structural motifs where alpha-helices interact to form dimers or higher order oligomeric states^13^. A repeating seven residues-long “heptad” motif with alternating hydrophobic (*h*) and polar (*p*) residues forming the general pattern *hpphppp* along the alpha-helices results in a core interface where the hydrophobic residues engage in knob-into-hole (KIH) interactions. The sequence-structure relationship of coiled-coils is well understood, and design principles have been established to create synthetic coiled-coils. Dimers are a specific example: it has been shown that isoleucine and leucine in positions *a* and *d* of a repeating heptad *abcdefg* (corresponding to the pattern *hpphppp*) greatly favor dimerization over other oligomeric states, whereas isoleucine at both positions favors trimers^14^. Moreover, the structure of coiled-coils can be parametrically modelled, which greatly accelerates the calculation of their physical properties, such as interaction strengths^15^. Parametric modelling entails the use of very few parameters (the radius, interface angle, and pitch of a coiled-coil) to describe the entire backbone structure. Parametric modelling, in combination with the BUDE (Bristol University Docking Engine)^16^ forcefield, has been successfully applied to predict the structures of natural and synthetic coiled-coils^17,18^.

In this study, we investigated the dimerization interface of MotB using parametric modelling to represent the MotB dimer as a dynamic coiled-coil. This simplification of the MotB structure enables the rapid calculation of the total energy of the system to discriminate between stable and unstable conformations. From these results, we rationally designed stator sequence variants to be tested *in vivo* and to better understand how MotB dimer stability may relate to overall conformational dynamics of the stator complex during its gating cycle.

In this study, we investigated the dimerization interface of MotB using parametric modelling to represent the MotB dimer as a dynamic coiled-coil. This simplification of the MotB structure enables the rapid calculation of the total energy of the system to discriminate between stable and unstable conformations. From these results, we rationally designed stator sequence variants to be tested *in vivo* and to better understand how MotB dimer stability may relate to the conformational dynamics of the stator complex during its gating cycle. By testing the *in vivo* motility MotB variants predicted to differ in stable and unstable conformational states, we aimed to identify which coiled-coil conformations were correlated with motility, and to inform the structural changes undertaken by *Ec*MotB and its extant homologs during their gating cycle.

## Results

We modelled the *Ec*MotB TM domain as a parallel coiled-coil based on the most recent structures of MotB from *B. subtilis* (PDBID: 6YSL), *C. sporogenes* (PDBID: 6YSF) and *C. jejuni* (PDBID: 6YKM)^7,8^ (Fig. 1). A set of ISAMBARD models of the 20-residue long *Ec*MotB peptide IAYA***D***FMTAMMAFFLVMWLI (corresponding to residues 28-47 of WT *Ec*MotB; the catalytic residue D32 is indicated in bold) with varying interface angles were aligned to the crystal structures to formally identify the “register” of the MotB coiled-coil, that is, what positions in the coiled-coil heptad each residue occupies. Based on RMSD calculations and the orientation of the catalytic aspartate (D32), we established that the 20-residue long MotB dimer stalk is best described as a coiled-coil starting at position *e*. In other words, residues A31, M34, M38, F41, and W45 occupy positions *a* and *d* along the coiled-coil, forming the core interface. These results agree with previous reports where mutation of these residues to cysteine led to crosslinking between the two MotB subunits, indicating proximity^19^. Further analysis using CCBuilder of the peptide IAYADFMTAMMAFFLVMWLI as a parallel dimeric coiled-coil starting at position *e* indicated that the five core residues could form KIH interactions.

**Figure 1.**
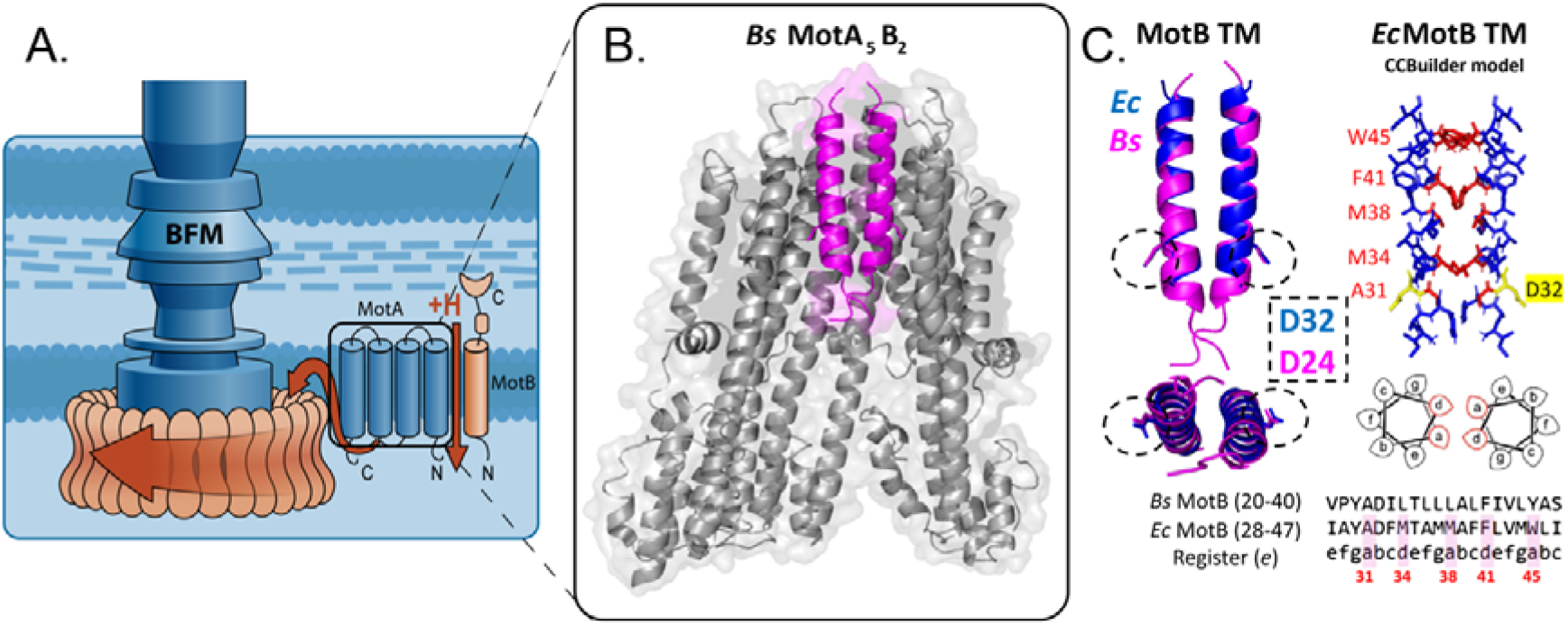
Modelling the transmembrane domain of MotB as a dimeric coiled-coil. A) The stator subunit of the bacterial flagellar motor (BFM) of *E. coli* (MotAB) drives the rotation of the large rotor complex by converting the proton motive force into mechanical torque. The stator unit MotA5MotB2 acts as an ion channel and each MotA has 4 transmembrane (TM) domains while MotB has a single TM domain and exists as a dimer in the centre of the complex. B) The crystal structure of *B. subtilis* MotA5B2[PDB 6YSL] (left) was used as the starting point for *E. coli* MotB TM modelling. The MotB dimer is shown in pink. C) A CCBuilder model of the homologous sequence from *E. coli* was generated (blue) and aligned to the reference *Bs*MotB dimer (purple). A coiled-coil model in the *e* register was selected based on the orientation of the catalytic aspartates D32 (*E*.*coli*) and D24 (*B. subtilis*) (middle, dotted circles in the side and bottom views of the TM domain). Residues A31, M34, M38, F41 and W45 were found to be at the coiled-coil interface (right), corresponding to positions *a* and *d* of the heptad repeat able to participate in knob-in-kole (KIH) interactions (labelled in red on the atomic model and heptad diagram underneath). An alignment of the two homologous amino acid sequences and the selected register is shown on the bottom-right, with *E*.*coli* residues at the *a* and *d* positions highlighted in red and numbered.

In light of these results, we decided to model MotB as a dimeric coiled to inform our *in vivo* mutagenesis experiments. To this end, rather than simply considering the most stable conformation calculated by ISAMBARD, we considered the conformational space of MotB by generating coiled-coil structural models with varying interface angles and radii and then, for each combination of the two parameter values, calculating the total energy of the system using the BUDE forcefield (hereafter, BUDE). This *in silico* conformational screen revealed the potential existence of wells (more stable configurations) and peaks (unstable configurations) across the energy landscape of the WT *Ec*MotB (Fig. 2A). Interestingly, the deepest stability well (#4, Fig. 2B) did not correspond to the configuration adopted in the cryo-EM structures (#1, Fig. 2B).

**Figure 2.**
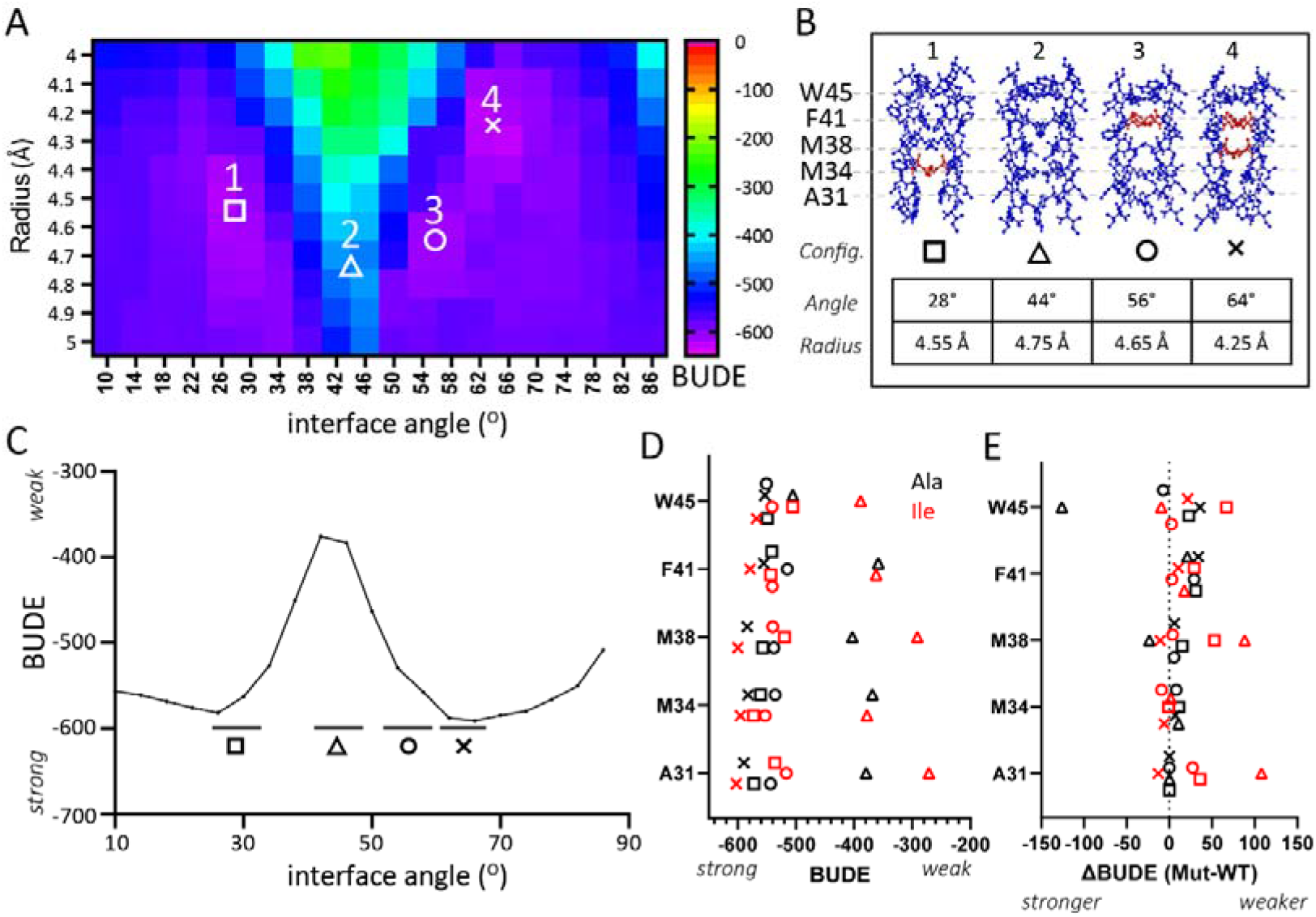
Characterization of the MotB TM coiled-coil interaction energy. (A) Free energy (BUDE) between MotB dimer coils represented as heatmap indicating BUDE value vs interface angle and radius. Four configurations are highlighted by symbols. (B) Atomic models of the MotB coiled-coil for four configurations (28°, 44°, 56° and 64°, identified by the square, triangle, circle and cross symbols respectively). The main amino acid residue responsible for KIH interactions are labelled in red. (C) 2D BUDE profile calculated from mean across ten radii values from (A). (D) predicted BUDE energy changes at each of the four configurations (symbols) after amino acid replacement at positions *a* and *d* of the MotB coil with amino acids alanine, (E) ΔBUDE (BUDE^mut^ – BUDE^wt^) for each of four configurations. The BUDE value calculated at interface angles 28°, 44° 56° and 64° is calculated as the average of the two neighbouring data points above the black horizontal bars. Substitutions are indicated by colour (Ala, black) or isoleucine (Ile, red).

Next, we considered strategic point mutation of coiled-coils to investigate if the stability of the MotB dimer would change after replacing the key *a* and *d* residues forming the core interface. We considered the design rules of synthetic coiled-coils and hypothesised that alanine mutations would systematically weaken the coiled-coil interaction by disrupting the hydrophobic KIH interactions, whereas isoleucine and leucine mutations would lead to more stable KIH interactions and thus a more stable coiled-coil (when isoleucine and leucine occupy positions *a* and *d*) or to steric clashes (when isoleucine occupies both positions *a* and *d*). Therefore, core residues were systematically mutated *in silico* to alanine, isoleucine, or leucine at positions *a* and *d* in over 200 permutations (see Method and Supplementary Table 1). In addition, we also considered the M38G mutation that appeared to disrupt KIH interactions in CCBuilder analyses. In total, a library of 221 sequence variants was generated *in silico*.

To test the above hypothesis *in silico*, we repeated the conformational analysis using ISAMBARD for each MotB variant modelled as a coiled-coil and compared the energy landscape of each sequence variant to the WT sequence. Different mutations resulted in a shift of the energy wells towards different interface angles and radii values and, in some extreme cases, to the absence of stability wells. The energy peaks corresponding to potential inaccessible conformations also differed. Altogether, the results indicate that the selected mutations in the core of the MotB dimerization interface could affect its accessible conformational space and, presumably, affect cell motility. Further to this analysis, the three-dimensional (3D) BUDE energy landscapes were collapsed into two-dimensional (2D) energy profiles by averaging the energy values for radii between 4 to 5 Å (Fig. 2C). This range of radii values in BUDE encompassed the radii observed in the reference structures (4.7 to 4.9 Å) and contained most of the energy wells and peaks of interest to compare sequence variants. In these dimensionally reduced datasets, we subsequently considered the interface angle parameter as the main variable affecting the energy of the coiled-coil system.

We next sought to select a subset of mutants to verify whether the differences in calculated energy profiles for MotB sequence variants correlated with BFM activity. Comparison of the 2D BUDE energy profiles suggested that single mutations could be sufficient to alter the accessible conformational space of MotB, and therefore affect cell motility (Fig. 2C, 2D). We selected 9 single mutants (SI Fig. 1) and an additional 10 double mutants and 4 triple mutants (SI Fig. 2) that showed varied energy profiles for *in vivo* experimentation. We restricted our *in vivo* testing to variants displaying alanine and/or isoleucine replacements.

We recombinantly expressed the selected sequence variants of MotB in an *E. coli* Δ*motAB* background strain and assessed the motility of bacteria on swim plates (Fig. 3A) and at a single cell level (Fig. 3B). Some of the single point mutations affected the size of swim rings (M34A, M34I, M38A, F41A) whereas all other mutations, including the selected double and triple mutants, abolished motility (Supplementary Fig. 3). Amongst the motile mutants, the first two had a comparable (M34A) or markedly positive (M34I) effect on motility, while M38A and F41A decreased the size of the swim rings (Fig.3A). Motility was confirmed at the single cell level when these four variants were tested in tunnel slides. Tethered cell assay measurement of cell speed revealed a significant reduction in cell speed for all the motile variants including M34I (p < 0.001, T test) (Fig. 3B). The M34I variant was also found to have a significant decrease in clockwise (CW) bias (p < 0.05) when compared to M34A indicating an increased propensity for counterclockwise (CCW) motor runs over tumbles (Supplementary Fig. 4), which may explain the increased size of swim rings for this variant in spite of the reduced motility. Interestingly, M34I was able to swim in higher density agar (0.4%) better than WT or any other variant, suggesting a change in torque generation in the mutant stator (SI Fig. 5).

**Figure 3.**
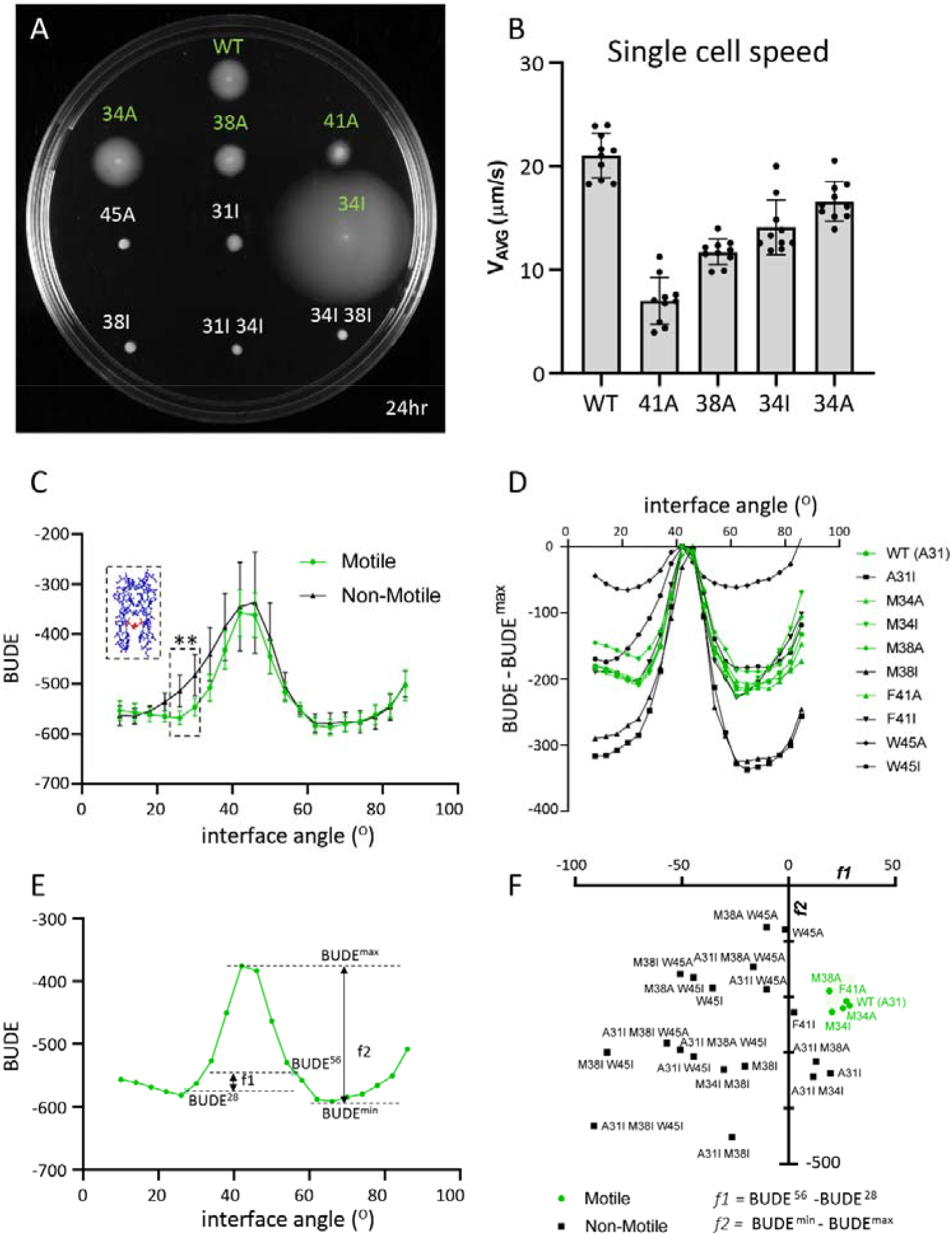
Testing coiled-coil variants in E.coli and parametrization of 2D BUDE profiles. A) Swim plate assay of *E. coli* ΔMotAB + pDB108 expressing WT MotA and either WT (top) or mutant MotB’s on LB agar 0.3% supplied with 0.1 % arabinose and Chloramphenicol after 24 hr incubation at 30’C. See supplementary figure 3 for all variants tested in this study. B) Single *E. coli* cell swimming speed measured in motility buffer. The 10 fastest cells for each mutant are plotted here. Bars indicate Mean speed (μm/s) + SD. C) Average 2D BUDE plots of motile (green) vs non-motile (black) MotB variants. The KIH present in MotB in the 28° configuration is highlighted in red on the atomic model in the dashed box. Bars indicate Mean + SD. D) Individual 2D BUDE plots of motile (green) vs non-motile (black) MotB variants, translated according to their respective BUDE^max^ values. E) The *f1* and *f2* parameters visualized on the 2D BUDE profile of *E. coli* WT MotB. F) Individual variants plotted on a graph (X = *f1* and Y = *f2*). The cluster of motile variants is highlighted in green.

**Figure 4.**
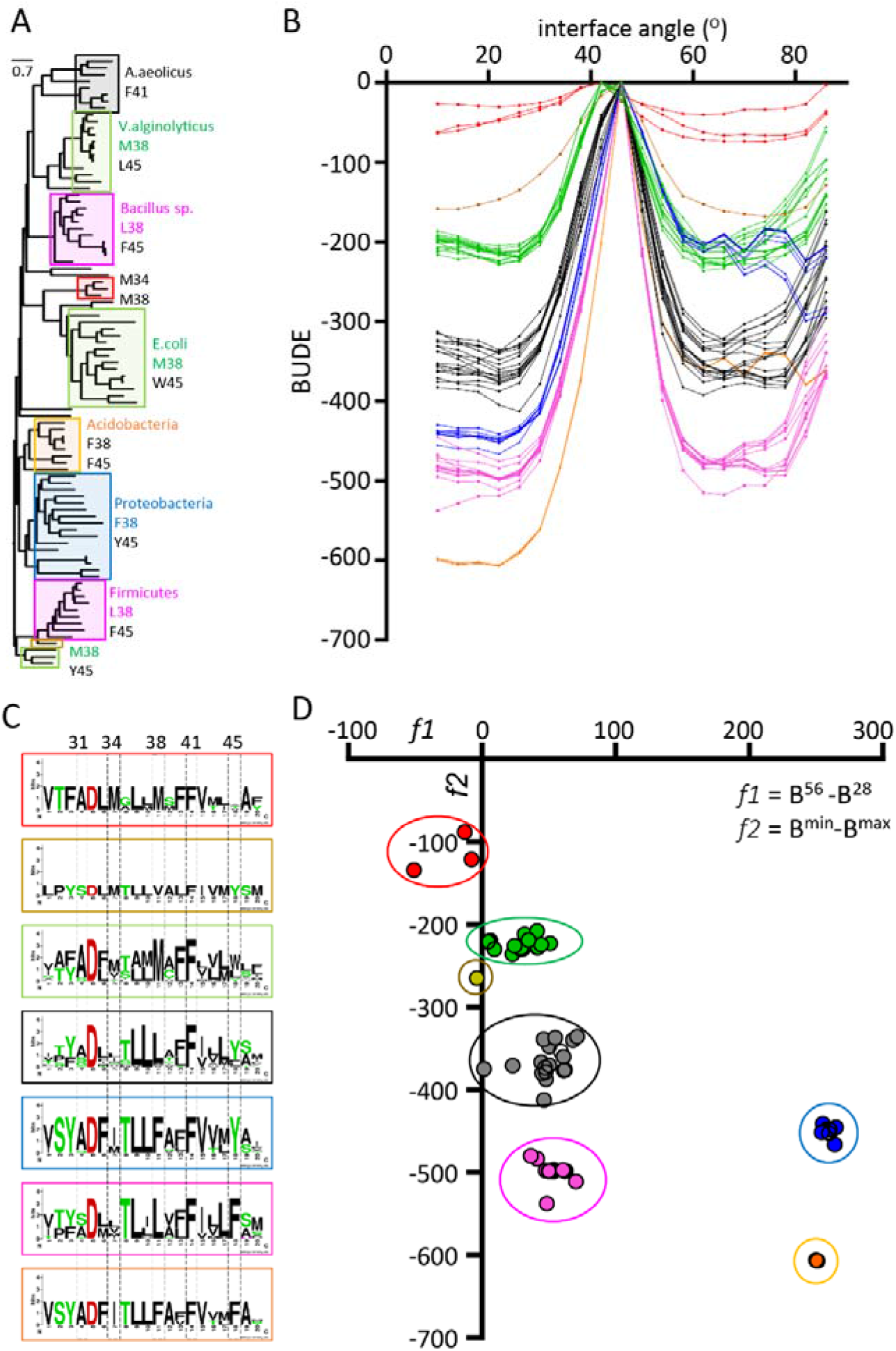
BUDE analysis of extant MotB sequences. A) Phylogeny of extant, full-length MotBs. Clades are labelled by colored boxes corresponding to the clusters highligted in B. A representative member of each clade, or representative bacterial phylum, are shown next to each box. The conserved amino acids indicative of each clade are shown next to each box and numbered according to *E*.*coli* residues. A detailed phylogeny is provided in SI Figure 10. B) 2D BUDE plots of extant WT MotB variants, translated by their BUDE^max^ values. C) Consensus logos for each plot in B. Each logo is inscribed in a box matching the colour of its respective plots in A. Knob-in-hole residues at positions homologous to *Ec*MotB 31, 34, 38, 41 and 45 are highlighted in vertical grey boxes. D) Clustered populations of extant MotB stators plotted on a graph (X = *f1* and Y = *f2*). The formulas used to calculate *f1* and *f2* for each homolog are provided below the X axis.

**Figure 5.**
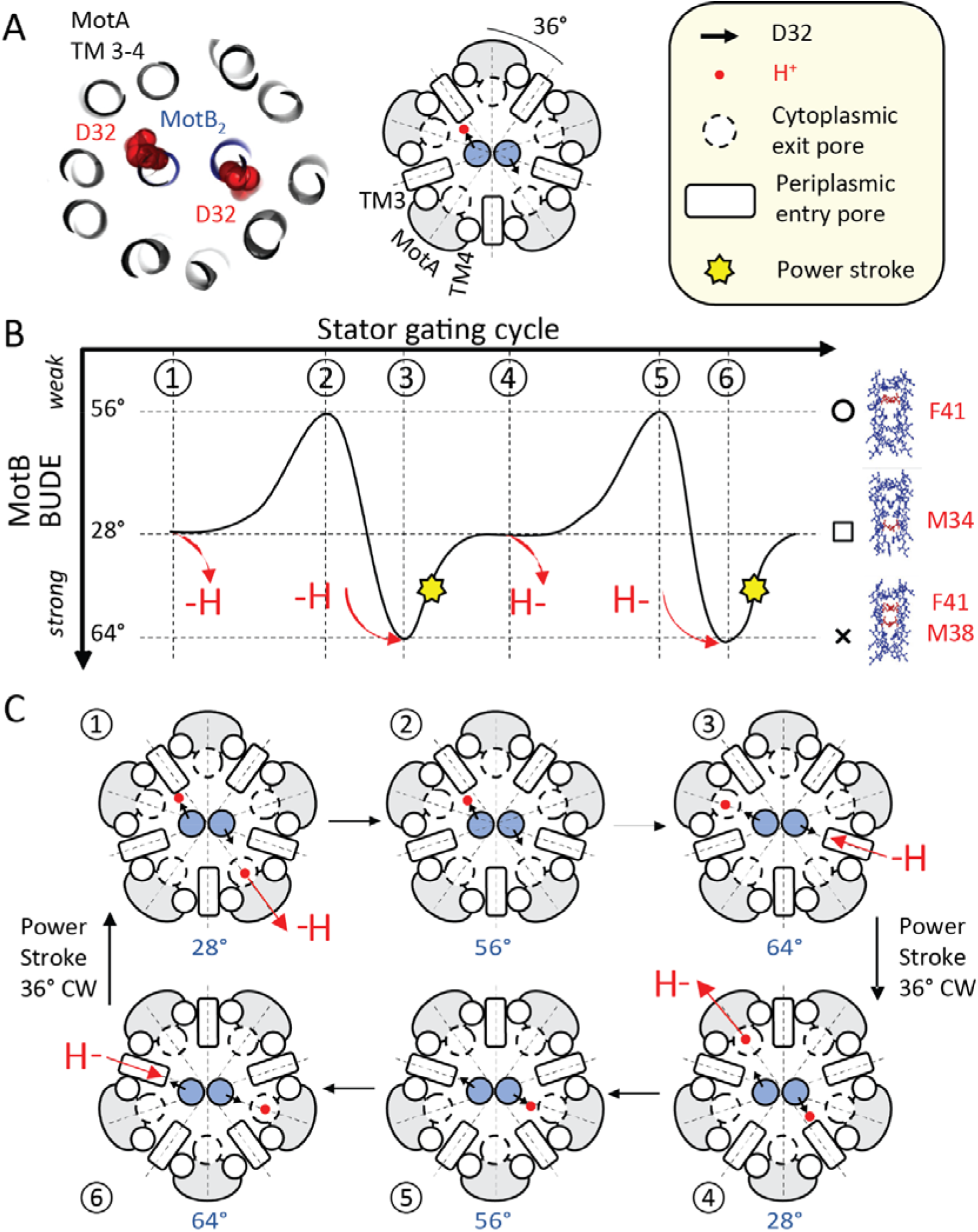
Rotary model showing the role of MotB coiled-coil conformations in the mechanochemical cycle of the stator complex. A) Section of the *Bs*MotAB [PDB 6YSL] structure highlighting the central MotB dimer (blue) and transmembrane helices TM3 and TM4 of MotA (grey) interfacing the MotB dimer. A sphere model (red) indicates the position and orientation of the catalytic aspartate in MotB. A cartoon representation of the crystal structure and legend are shown on the right. The dashed lines below separate the stator cross-section in ten identical 36° sectors. B) MotB TM BUDE fluctuations between three configurations during the gating cycle of the stator. The red arrows indicate binding or unbinding of a proton at the left or right aspartate (H-or -H, respectively). A yellow star indicates the power stroke and 36° clockwise rotation of the stator. Configuration symbols and representative KIH interactions are also shown on the right of the model. C) Six stages of the mechanochemical cycle model described in B. A proton is transferred into the cytoplasm through an exit pore immediately after a power stroke (1). A change in the MotB TM configuration follows, which rotates 36° counterclockwise from the 28° (1) to the 64° configuration (3), via the 56° configuration (2). Once in the 64° configuration, the aspartate on the left subunit engages MotA at the TM3-4 interface in proximity of the cytoplasmic exit pore (3). A proton then enters from the opposite periplasmic entry pore to bind the unprotonated aspartate causing a conformational change of the coiled-coil and the coordinated rotation of the MotB dimer and MotA pentamer by 36° clockwise (a power stroke). This rotation restores the coiled-coil to the 28° configuration and the engaged ion is released into the cytoplasm. The cycle repeats at the previously disengaged aspartate in (4), (5) and (6) eventually resulting in a new power stroke and the return to stage (1).

A comparison of the mean 2D BUDE profiles of all motile and non-motile variants suggested that the stability of the coiled-coil around the interface angle of 28°, where M34 plays a key role in KIH interaction, was crucial for function (Fig. 3C). The influence on the total energy from mutation at that site, adjacent to the catalytic aspartate residue (D32), appears to be correlated with a motile phenotype. When comparing the 2D BUDE profiles of each MotB variant tested *in vivo*, we noticed a similarity in the shapes of the BUDE profiles of the motile variants, whereas the profiles of the non-motile variants diverged substantially (Fig. 3D). Aligning the profiles with respect to their peak values (BUDE^max^) revealed additional differences between motile and non-motile profiles than the strength of interaction at the 28° configuration alone. The BUDE gap between the minimum value of each profile and the peak corresponding to the 44° configuration appeared to be similar across the motile variants.

Further comparison of the 2D profiles of all point mutants suggested a pattern of specific interface angles, or configurations, associated with motility (SI Fig. 6). All the *Ec*MotB variants found to be motile had the total energies associated with the 66° (cross), 28° (square), 56° (circle) and 44° (triangle) configurations in ascending order of magnitude. The exceptions to this pattern were the single mutants A31I, F41I and M38G, which featured destabilized 44°, 28°, and 28° configurations, respectively, and the double mutants A31I M34I and A31I M38A which featured a destabilized 44° configuration. This suggested that the relative stability of the 28°, 56° and 64° configurations play a role in determining the function of MotB and that this pattern may reflect the order of configurations undertaken by WT MotB during its gating.

We captured the features associated with motility variants by defining two metrics, *f1* and *f2* (Fig. 2E), where *f1* = BUDE^56°^ -BUDE^28°^ and *f2* = BUDE^min^ – BUDE^max^ (Fig. 3E). The first metric, *f1*, is the BUDE energy gap at an interface angle of 28°, associated with motile variants (Fig 3C), and the reference interface angle of 56°, predicted to form a KIH interaction with the conserved residue F41 (Fig. 2B, ‘circle’ config). Residue F41 appeared to have the more consistent value across the profiles of all variants (SI Fig. 7). With the *f2* metric we measured the energy gap between the minimum and maximum BUDE values across the profile, or the energy barrier separating the energy wells. Separating all variants on the basis of these two metrics revealed a cluster of optimal *f1* and *f2* values for the motile variants of *E. coli* MotB (Fig. 3F). These results suggest that a combination of energetic parameters contained within the 2D profiles could be descriptive of functional *Ec*MotB variants and that the relative stability of each configuration needs to be maintained to operate the *E. coli* stator.

We also searched for other metrics that could cluster and separate motile and non-motile sequences to facilitate the prediction of motile variants of *Ec*MotB from their 2D profile models. We defined *f3 =* [(BUDE^56°^ -BUDE^28°^) -(ΔBUDE^56°^ -ΔBUDE^28°^)] and observed that this metric was correlated with motility only when yielding a positive value, where ΔBUDE is the difference in BUDE energy between the mutant and WT MotB at the given interface angle. Additionally, we defined *f4 =* (BUDE^min^ -BUDE^max^ -BUDE^max^) = (BUDE^min^ −2*BUDE^max^). This considered not only the range (the difference between min and max) but also the maximum value as an indicator of the magnitude of the BUDE value (allowing, e.g., to distinguish between two samples with range of 200, but with a BUDE^max^ of −500 or −400 respectively). This quantity, *f4*, was found to correlate with motility when positive. We observed that both *f3* and *f4* were positive only in the functional variants of *E. coli* MotB studied here (Supplementary Fig. 8).

Finally, we explored a larger set of homologous MotB sequences to determine whether our observations were applicable to other extant motile bacteria. We generated a 91 member phylogeny of MotB using RAxML^20^ across diverse clades (Fig. 4A, SI Fig. 9, 10) and carried out the same BUDE analysis on 70 representative MotB sequences across the phylogeny (Supplementary Fig. 9A, 10) to produce a 2D BUDE energy profile for each of the 20 residues-long TM domain modelled as a coiled-coil (SI Fig. 9B). To compare the energy profiles to each other and to the phylogenetic clustering, we translated each BUDE profile relative to its BUDE^max^ value (Fig. 4B). This analysis of the energy profiles distinguished 7 distinct groups (Fig. 4C) that matched the phylogenetic clades (Fig. 4A), and each displaying one or more conserved amino acid residues amongst the representative positions *a* and *d* of their heptad repeats, evidenced by their consensus logos (Fig. 4C). These groups clustered around specific values of the *f1* and *f2* metrics (Fig. 4D). Out of these 70 sequences, 66 were found to have *f1* > 0, while only a subset had *f2* > 0 (SI Fig. 9C). Clusters were separated by the identity of their residue 38 (position *a*) and shared the conserved D32 and F41 (*d*) residues. Positions 31 (*a*), 34 (*d*) and 45 (*a*) also displayed unique residues in specific clusters (Fig. 4A, 4C).

## Discussion

In this study we constructed several MotB stator variants based on *in silico* designs following coiled-coil design principles and measured their capacity to drive swimming *in vivo* in *E. coli*. Because the residues targeted by mutagenesis were at the hydrophobic interface of the coiled-coil structure, we assumed that the effects of amino acid replacement would be confined to MotB and its dimerization and thus restricted our analyses to symmetric configurations of the MotB dimer. In other words, we assumed that movement (i.e. rotation) of one MotB subunit was always accompanied by the same movement of the second subunit. We sought to examine how motility of the overall stator complex was disrupted when assembly of the coiled-coil via knobs-in-holes (KIH) was altered. Altogether, we focused on variations of the interface angle between the two MotB alpha-helices, which correspond to the rotation of the alpha-helices relative to each other. This type of rotation is known as a register shift, as the interface angle is related to the coiled-coil register, and register shifts have been shown elsewhere to underpin biological processes such as dynein recruitment^21,22^ and the activity of proteasomal ATPase^23^.

We tested a mutant with the intent to disrupt motility (M38G), by disrupting the KIH interactions between the coils. A gain-of-function variant (M34I), designed to have a stronger coiled-coil interaction, demonstrated that coiled-coil stability could be linked to improved motility, but we also observed that “tightening” the coiled-coil at the wrong knob could disrupt function entirely. For example, mutating the central knob towards a new type of KIH interaction (M38I mutant) was predicted to stabilize only the 64° conformation. This mutation yielded a non-motile variant. In contrast, energy calculations indicated the M34I mutation could increase dimer stability for all analysed coiled coil configurations, including the 64° confirmation, and yet this mutation yielded a motile variant. The reason for improved swimming of this variant on swim plates remains unclear because the motor performance appeared to be reduced compared to WT in single-cell assays (both in tethered and free-swimming assays). In single-cell assays, M34I was indistinguishable from the M34A variant, while M34A displayed WT phenotype on swim plates in contrast to M34I.

Swimming of M34I, WT, M34A, M38A and F41A correlated with shared properties of their BUDE energy profiles. Although a bulk comparison of our motile and non-motile variants showed a difference in their BUDE magnitude at a specific configuration on the profile (the 28° configuration) we sought to identify other features in the BUDE profile to use for classifying motile and non-motile classes. We defined two metrics, *f1* and *f2*, to differentiate between the two classes and the effect of each of point mutant on motility. In this framework, M38A maintains the energy profile shape of the WT; M38I reduces BUDE^max^ and abolishes function; F41A destabilizes the 56° configuration, maintaining a positive value for *f1* and therefore a functional stator, while F41I stabilizes it and abolishes function by lowering *f1* below zero; W45A increases BUDE^max^ beyond a functional level and W45I destabilizes the 28°configurations, thus lowering *f1* to a negative value. A31I reduces BUDE^max^ compared to WT and abolishes function.

Using *f1* as an indicator of functionality in a stator implies that the interaction is dominated by the knobs at position 34 and the conserved residue F41. We thus speculate that a transition between these two configurations is essential for the function of the stator. Previous work involving chemical cross-linking between MotB coiled-coil residues suggested that the TM structure is flexible and able to exchange the residues at the CC interface during a rotation of ∼36°^19^. In line with those considerations, we found that BUDE profile features at all four configurations (28°, 44°, 56° and 64° configurations in *Ec*MotB) were predictive of stator functionality, that is, motility on swim plates.

The *f1* and *f2* parameters were useful also in describing and clustering extant MotBs. Indeed, it may be that these clusters represent families of MotB homologs that are related in their functional characteristics. Notably, *V. alginolyticus* PomB and *E. coli* MotB, of which functional chimeras exist^24^, belong to the same cluster (green, Fig. 4). These two stators are known to differ in their power source (Na^+^ and H^+^ respectively) and differ in the identity of their homologous residue 45 (L45 for Na^+^, W45 for H^+^) while showing similar 2D profiles. Another notable cluster is the one shared by *B. subtilis, C. sporogenes* and *C. jejuni* (black, Fig. 3). These three species are sources of the most detailed cryo-EM structures of the MotA_5_B_2_ stator complex to date. It is possible that these complexes were more stable and thus easier to purify and resolve due to stronger MotB coil interaction than the *E. coli* or *V. alginolyticus* homologs.

We noted a linear relationship between the hydrophobicity of residue 38 with the BUDE^max^ magnitude. MotB clusters displaying F, L or M at position 38 had a proportionally high BUDE^max^ peak in their profiles, with F38 being the highest and M38 the lowest (SI Fig. 9B). The height of this peak could be related to the presence of stable intermediate conformations available during the transition between the two most stable configurations at 28° and 64°.

Elsewhere we have extensively examined large phylogenies of MotB^25^. Here we calculated a small phylogeny that was representative of wider flagellar diversity to compare how energetics clustered across a model of evolutionary history. Our MotB classes obtained from clustering similar energetic profiles closely overlap clades from our phylogeny obtained using traditional tree-building approach (Fig. 3A). This suggests that conservation of sequence identity is closely linked to conservation of functional traits, and indeed it is notable that BUDE free energy profiles calculated across only the 20 transmembrane residues so naturally fell into clusters that matched those generated by sequence similarity from phylogenetic consideration across the full length MotB. Together, these results show that cluster-based comparison of BUDE free energies can recapitulate phylogenetic clades and shows that there is evolutionary evidence for conservation of distinct stable states. This clustering based on MotB-MotB dimer stability could be further examined in greater detail, for example, with regard to species that have specific high-torque operating environments (e.g., *Spirochaetes* sp.). These species typically have large rotors to generate larger torques^5,26–28^, but there could be signatures of conservation of increased stator stability elsewhere that may correlate with high-torque or viscous operating conditions.

Based on the model proposed by Rieu *et al*.^10^, we describe a model of mechanochemical gating that includes the conformational changes in the MotB TM domain studied here (Fig.5). Our model assumes that the rotation of the MotA pentamer can occur only if both catalytic aspartates in MotB are protonated. We propose that a single protonated aspartate, engaging MotA with MotB, couples the synchronized rotation of both MotA_5_ and MotB_2_ by 36° clockwise, and is then released into the cytoplasm after the entry of a second proton binding onto the other disengaged aspartate. Our model also assumes that the binding of a proton to the disengaged aspartate initiates the power stroke, which then occurs as both aspartates are protonated as postulated by Santiveri et al.^7,10^ The newly protonated site triggers a conformational change at the MotA-MotB interface that promotes the engagement of the protonated aspartate with MotA at the second catalytic aspartate, the cytoplasmic pore exit site, as the complex rotates 36° clockwise.

In the Oster *et al*.^4^ model, a ‘kink’ occurs in TM3 of MotA upon H^+^ binding to the aspartate and when MotB engages with TM3/4 of MotA this results in a swivelling motion of the TM3 of MotA that is the basis of the power stroke mechanism. Our data suggests that MotB might be coupled with the movement of MotA during its swivelling motion (a CW rotation) and switch between at least two of the stable configurations studied here, the 28° and 64° ones. Our model takes into account the 5:2 stoichiometry detected in crystal structures and considers only one catalytic site at the time to be participating in a power stroke. The concerted CW rotation of the MotB TMs during the power stroke facilitates the alternating engagement of MotA subunits separated by 2/5^th^ of a revolution, while maintaining MotB always in a parallel coiled coil state and without the need for vertical displacement of the MotB coils during the proton release stage. The activation of individual MotA subunits during the power stroke is reminiscent of the model proposed by Xing *et al*. but involving the alternating translocation of protons across the stator^29^.

Whilst we cannot determine the exact motions of the MotB TM dimer during the switch between its two most stable configurations, we suspect that the relative stability of the intermediate configurations (like the 56° one) is important for the completion of a 36° rotation of the interface angle of MotB. It is also possible for other coiled-coil parameters (pitch and radius) to change during this rotation which may stabilise the coiled coil at configurations considered inaccessible (like the 44° configuration). Mutations that further stabilise certain configurations, for example a more hydrophobic KIH at position 38 of the *Ec*MotB, would increase the height of the BUDE^max^ peak (Fig.4). This might interfere with the establishment of these intermediate conformations which need to be crossed during the rotation from the 28° to the 64° configuration.

The function of the flagellar stator complex clearly depends not only on the pore lining residues between MotA and MotB and torque-generating interactions between MotA and FliG^9,30^. Our overall conclusion is that it also depends on the degree of interaction between the two TM domains of MotB that assemble as a coiled-coil at the core of the stator and the conformational states the dimer can access. The residues responsible for knob-in-hole interactions at the dimerization interface of *E. coli* MotB, and possibly other extant homologs, appear to tolerate onlyl those mutations which do not shift their coiled-coil energetics too far from wild-type. Consideration of these energetics will improve our ability to engineer mutant MotB subunits, chimeric stators, or synthetic combinations of multiple flagellar motor proteins.

## Materials and Methods

### Structural modelling of coiled-coils

The *B. subtilis* MotA_5_B_2_ structure (PDB:6YSL)^8^ was used as a reference to model the transmembrane domain of *E. coli* MotB as coiled-coil. We used ISAMBARD to create a specification whereby a parallel dimeric coiled-coil backbone structure could be generated from three parameters: the interface angle, the radius, and the pitch. The coiled-coil backbone structure for the amino acid sequence IAYADFMTAMMAFFLVMWLI (corresponding to the *E. coli* MotB residues 28-47) was optimized using the genetic algorithm evolutionary optimizer from ISAMBARD and SCWRL 4.0 for the optimization of side chain orientations to minimize the total energy of the system as calculated by the BUDE force field. Optimizations were performed using seven different initial interface angle values which corresponded to the seven possible registers of a coiled-coil. The seven optimized coiled-coil models were aligned to the resolved structure of the MotB dimer stalk in PDB structure 6YSL. The alignment revealed that the best initial interface angle that resulted in a good structural match after optimization to the MotB stalk was 51.4°, which corresponds to position *e* for the first residue in the amino acid sequence. The optimized parameter values were 4.54 Å radius, 232.03 Å pitch, and 74.85° interaction angle.

### Calculation of the energy landscapes of the MotB coiled-coils

We used in-house Python scripts to generate a list of 221 sequence variants of the amino acid sequence IAY**A**DF**M**TAM**M**AF**F**LVM**W**LI where residues A31, M34, M38, F41, W45 were systematically mutated to alanine in every permutation; separately the sequence was mutated with isoleucine (for residues A31, M38, and W45), asparagine (for residue M38), and leucine (for residues M34 and F41) in every permutation. These mutations follow coiled-coil design principles to weaken or strengthen dimeric coiled-coils. Finally, additional MotB mutants were added to the list (SI Table 1). For each sequence variant, we used ISAMBARD to generate coiled-coil backbone structural models with radii values ranging from 4 to 6 Å (in 0.1 Å increments) and interface angle values ranging from 10 to 86° (in 4° increments). The pitch was kept constant to a value of to 150 Å. These ranges of values encompass the optimized parameters that best matched the reference structures. The sidechain orientations were optimised using SCWRL 4.0 as part of the ISAMBARD script. For each conformational variant, the total energy of the system was calculated using the BUDE forcefield. The three-dimensional data sets (BUDE total energy as a function of interface angle and radius) were plotted as heatmaps using in-house Python scripts and Matplotlib library. Two-dimensional datasets were generated by integrating the total energy values for a subset of the radii values (4 to 5 Å) which contained the stability wells and instability peaks of interest. The integrated total energy values were plotted as a function of the interface angles. In some analyses, we focused on four regions in the three-dimensional dataset for comparison of the sequence variants (centred on list combinations of radii and interface angles) and a representative energy value for each of the four regions was calculated by averaging two neighbouring points from the two-dimensional dataset (see Fig. 2C). For the identification of knob-into-hole interactions, MotB was modelled as a dimeric coiled-coil starting at register *e* using CCBuilder 2.0, which evaluates the presence of knob-into-holes using SOCKET^31^.

### *E. coli* strains, plasmids, and culture media

*E. coli* strain RP437 Δ*motAB* was prepared using the No-Scar method^32^. MotA and MotB were expressed from plasmid pDB108 (Cm^+^) after induction with 0.1 mM Arabinose. FliC-sticky *E. coli* RP437 cells (Δ*motAB* Δ*cheY* Δ*pilA fliC*^*st*^)^33^ were used to perform the tethered cell assay experiments. Liquid cell culturing was done using LB broth (85 mM NaCl, 0.5% yeast extract, and 1% Bacto tryptone). Point mutations in *motB* on the pDB108 plasmid were generated using the QuikChange^™^ technique and verified by Sanger sequencing. A list of primers used is provided in Supplementary Table 2. Cells were cultured on agar plates composed of LB broth and 1% Bacto agar (BD Biosciences, U.S.A.). Swim plate cultures were performed on the same substrates adjusted for agar content (0.3% Bacto agar).

### Single cell swimming assay and tethered cell assay

Free-swimming cells were grown overnight in TB buffer (85 mM NaCl, 1% Bacto tryptone) at 30°C to OD_600_ of ∼0.5 then washed three times in 1 ml MB buffer (85 mM NaCl and 10 mM KPi, pH=7.0) before resuspension in 500 μl of MB buffer and imaging in a tunnel slide. The tethered cell assay was performed as previously described^34^. Briefly, 1 ml of cells [optical density at 600 nm (OD600) = 0.5] grown in TB buffer was sheared by passing the cell suspension through a 26-gauge syringe needle 30 times. These cells were then washed three times in 1 ml MB and lastly resuspended in 500 μl of the same buffer. Then, 20 μL of suspension was loaded into a tunnel slide prefilled with MB buffer and incubated for 10 min at room temperature. The unbound cells were then removed from the tunnel slide by washing with a total of 200 μl of MB buffer (∼10 times the tunnel slide volume) and then loaded onto a phase contrast microscope for video recording. Time-lapse videos were recorded at 40x magnification on a phase contrast microscope (Nikon). Time-lapse videos were collected using a camera (Chameleon3 CM3, Point Grey Research) recording 20-s-long videos at 20 frames/s. Experiments involving single-cell tracking were recorded at 60 frames/s. A custom LabView software^35^ was used as previously reported to estimate specific rotational parameters of the tethered cells such as rotation frequency (speed), clockwise and counterclockwise bias, switching frequency, and speed of swimming cells.

## REFERENCES

1. Nakamura, S., and Minamino, T. (2019). Flagella-driven motility of bacteria. Biomolecules 9, 279. 10.3390/biom9070279.

2. Mondino, S., San Martin, F., and Buschiazzo, A. (2022). 3D cryo-EM imaging of bacterial flagella: Novel structural and mechanistic insights into cell motility. Journal of Biological Chemistry 298, 102105. 10.1016/j.jbc.2022.102105.

3. Nirody, J.A., Sun, Y.-R., and Lo, C.-J. (2017). The biophysicist’s guide to the bacterial flagellar motor. Advances in Physics: X 2, 324–343. 10.1080/23746149.2017.1289120.

4. Mandadapu, K.K., Nirody, J.A., Berry, R.M., and Oster, G. (2015). Mechanics of torque generation in the bacterial flagellar motor. Proc Natl Acad Sci USA 112, E4381–E4389. 10.1073/pnas.1501734112.

5. Guo, S., and Liu, J. (2022). The Bacterial Flagellar Motor: Insights Into Torque Generation, Rotational Switching, and Mechanosensing. Frontiers in Microbiology 13.

6. Hu, H., Santiveri, M., Wadhwa, N., Berg, H.C., Erhardt, M., and Taylor, N.M.I. (2022). Structural basis of torque generation in the bi-directional bacterial flagellar motor. Trends in Biochemical Sciences 47, 160–172. 10.1016/j.tibs.2021.06.005.

7. Santiveri, M., Roa-Eguiara, A., Kühne, C., Wadhwa, N., Hu, H., Berg, H.C., Erhardt, M., and Taylor, N.M.I. (2020). Structure and Function of Stator Units of the Bacterial Flagellar Motor. Cell 183, 244–257.e16. 10.1016/j.cell.2020.08.016.

8. Deme, J.C., Johnson, S., Vickery, O., Muellbauer, A., Monkhouse, H., Griffiths, T., James, R.H., Berks, B.C., Coulton, J.W., Stansfeld, P.J., et al. (2020). Structures of the stator complex that drives rotation of the bacterial flagellum. Nature Microbiology, 1–12. 10.1038/s41564-020-0788-8.

9. Hu, H., Popp, P.F., Santiveri, M., Roa-Eguiara, A., Yan, Y., Liu, Z., Wadhwa, N., Wang, Y., Erhardt, M., and Taylor, N.M.I. (2022). Mechanisms of ion selectivity and rotor coupling in the bacterial flagellar sodium-driven stator unit. 2022.11.25.517900. 10.1101/2022.11.25.517900.

10. Rieu, M., Krutyholowa, R., Taylor, N.M.I., and Berry, R.M. (2022). A new class of biological ion-driven rotary molecular motors with 5:2 symmetry. Frontiers in Microbiology 13.

11. Sharp, L.L., Zhou, J., and Blair, D.F. (1995). Tryptophan-scanning mutagenesis of MotB, an integral membrane protein essential for flagellar rotation in Escherichia coli. Biochemistry 34, 9166–9171. 10.1021/bi00028a028.

12. Zhang, X.Y.-Z., Goemaere, E.L., Seddiki, N., Célia, H., Gavioli, M., Cascales, E., and Lloubes, R. (2011). Mapping the interactions between Escherichia coli TolQ transmembrane segments. J. Biol. Chem. 286, 11756–11764. 10.1074/jbc.M110.192773.

13. Lupas, A.N., and Bassler, J. (2017). Coiled Coils – A Model System for the 21st Century. Trends in Biochemical Sciences 42, 130–140. 10.1016/j.tibs.2016.10.007.

14. Fletcher, J.M., Boyle, A.L., Bruning, M., Bartlett, G.J., Vincent, T.L., Zaccai, N.R., Armstrong, C.T., Bromley, E.H.C., Booth, P.J., Brady, R.L., et al. (2012). A Basis Set of de Novo Coiled-Coil Peptide Oligomers for Rational Protein Design and Synthetic Biology. ACS Synth. Biol. 1, 240–250. 10.1021/sb300028q.

15. Crick, F.H.C. (1953). The Fourier transform of a coiled-coil. Acta Cryst 6, 685–689. 10.1107/S0365110X53001952.

16. McIntosh-Smith, S., Price, J., Sessions, R.B., and Ibarra, A.A. (2015). High performance in silico virtual drug screening on many-core processors. The International Journal of High Performance Computing Applications 29, 119–134. 10.1177/1094342014528252.

17. Wood, C.W., Bruning, M., Ibarra, A.Á., Bartlett, G.J., Thomson, A.R., Sessions, R.B., Brady, R.L., and Woolfson, D.N. (2014). CCBuilder: an interactive web-based tool for building, designing and assessing coiled-coil protein assemblies. Bioinformatics 30, 3029–3035. 10.1093/bioinformatics/btu502.

18. Wood, C.W., and Woolfson, D.N. (2018). CCBuilder 2.0: Powerful and accessible coiled-coil modeling: Powerful and Accessible Coiled-Coil Modeling. Protein Science 27, 103–111. 10.1002/pro.3279.

19. Braun, T.F., and Blair, D.F. (2001). Targeted disulfide cross-linking of the MotB protein of Escherichia coli: evidence for two H(+) channels in the stator Complex. Biochemistry 40, 13051–13059. 10.1021/bi011264g.

20. Stamatakis, A. (2014). RAxML version 8: a tool for phylogenetic analysis and post-analysis of large phylogenies. Bioinformatics 30, 1312–1313. 10.1093/bioinformatics/btu033.

21. Choi, J., Park, H., and Seok, C. (2011). How does a registry change in dynein’s coiled-coil stalk drive binding of dynein to microtubules. Biochemistry 50, 7629–7636. 10.1021/bi200834k.

22. Noell, C.R., Loh, J.Y., Debler, E.W., Loftus, K.M., Cui, H., Russ, B.B., Zhang, K., Goyal, P., and Solmaz, S.R. (2019). Role of Coiled-Coil Registry Shifts in the Activation of Human Bicaudal D2 for Dynein Recruitment upon Cargo Binding. J. Phys. Chem. Lett. 10, 4362–4367. 10.1021/acs.jpclett.9b01865.

23. Snoberger, A., Brettrager, E.J., and Smith, D.M. (2018). Conformational switching in the coiled-coil domains of a proteasomal ATPase regulates substrate processing. Nat Commun 9, 2374. 10.1038/s41467-018-04731-6.

24. Nishino, Y., Onoue, Y., Kojima, S., and Homma, M. (2015). Functional chimeras of flagellar stator proteins between E. coli MotB and Vibrio PomB at the periplasmic region in Vibrio or E. coli. Microbiologyopen 4, 323–331. 10.1002/mbo3.240.

25. Islam, M.I., Lin, A., Lai, Y.-W., Matzke, N.J., and Baker, M.A.B. (2020). Ancestral Sequence Reconstructions of MotB Are Proton-Motile and Require MotA for Motility. Front. Microbiol. 11. 10.3389/fmicb.2020.625837.

26. Beeby, M., Ferreira, J.L., Tripp, P., Albers, S.-V., and Mitchell, D.R. (2020). Propulsive nanomachines: the convergent evolution of archaella, flagella and cilia. FEMS Microbiology Reviews 44, 253–304. 10.1093/femsre/fuaa006.

27. Beeby, M., Ribardo, D.A., Brennan, C.A., Ruby, E.G., Jensen, G.J., Hendrixson, D.R., and Hultgren, S.J. (2016). Diverse high-torque bacterial flagellar motors assemble wider stator rings using a conserved protein scaffold. Proceedings of the National Academy of Sciences of the United States of America 113, E1917–E1926. 10.1073/pnas.1518952113.

28. Carroll, B.L., and Liu, J. (2020). Structural Conservation and Adaptation of the Bacterial Flagella Motor. Biomolecules 10, 1492. 10.3390/biom10111492.

29. Xing, J., Bai, F., Berry, R., and Oster, G. (2006). Torque–speed relationship of the bacterial flagellar motor. Proceedings of the National Academy of Sciences of the United States of America 103, 1260–1265. 10.1073/pnas.0507959103.

30. Islam, M.I., Ridone, P., Lin, A., Michie, K.A., Matzke, N.J., Hochberg, G., and Baker, M.A. (2022). Ancestral reconstruction of the MotA stator subunit reveals that conserved residues far from the pore are required to drive flagellar motility. 2022.10.17.512626. 10.1101/2022.10.17.512626.

31. Walshaw, J., and Woolfson, D.N. (2001). SOCKET: a program for identifying and analysing coiled-coil motifs within protein structures11Edited by J. Thornton. Journal of Molecular Biology 307, 1427–1450. 10.1006/jmbi.2001.4545.

32. Reisch, C.R., and Prather, K.L.J. (2015). The no-SCAR (Scarless Cas9 Assisted Recombineering) system for genome editing in Escherichia coli. Sci Rep 5, 15096. 10.1038/srep15096.

33. Sowa, Y., Rowe, A.D., Leake, M.C., Yakushi, T., Homma, M., Ishijima, A., and Berry, R.M. (2005). Direct observation of steps in rotation of the bacterial flagellar motor. Nature 437, 916–919. 10.1038/nature04003.

34. Islam, M.I., Bae, J.H., Ishida, T., Ridone, P., Lin, J., Kelso, M.J., Sowa, Y., Buckley, B.J., and Baker, M. a. B. (2021). Novel Amiloride Derivatives That Inhibit Bacterial Motility across Multiple Strains and Stator Types. J Bacteriol 203, e0036721. 10.1128/JB.00367-21.

35. Ishida, T., Ito, R., Clark, J., Matzke, N.J., Sowa, Y., and Baker, M.A.B. (2019). Sodium-powered stators of the bacterial flagellar motor can generate torque in the presence of phenamil with mutations near the peptidoglycan-binding region. Molecular Microbiology 111, 1689–1699. https://doi.org/10.1111/mmi.14246.

