## Supplementary Figures 1-10, Supplementary Tables 1&2. for "Tuning the stator subunit of the flagellar motor with coiled-coil engineering"

**Supplementary Material**

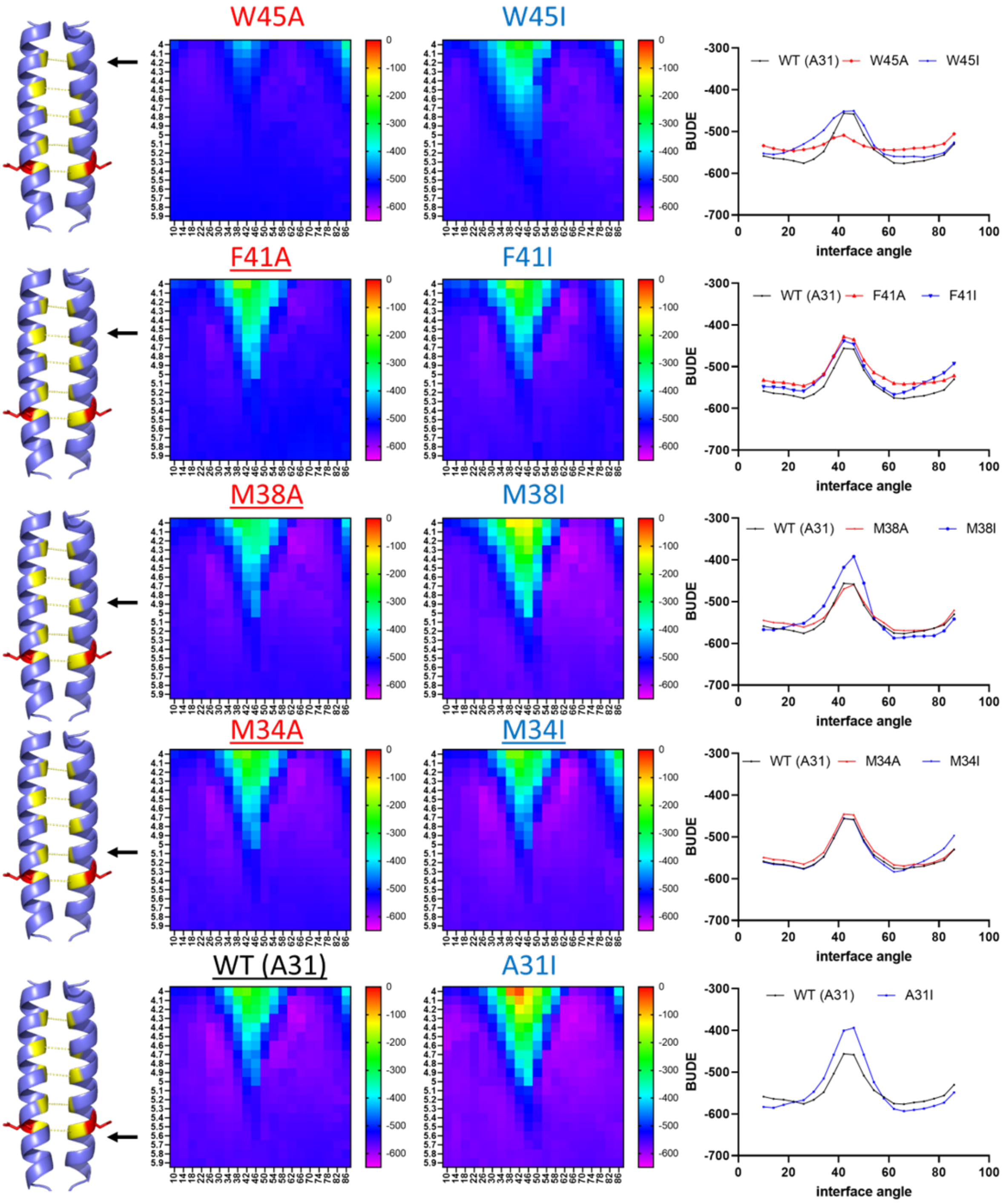

**Supplementary Figure 1. BUDE modelling of MotB variants.**

Schematic representation of the position of the amino acid residue being mutated to alanine or isoleucine along the TM domain of MotB (left). Heatmaps (centre) and 2D BUDE profiles (right) of each single point mutant tested in this study.

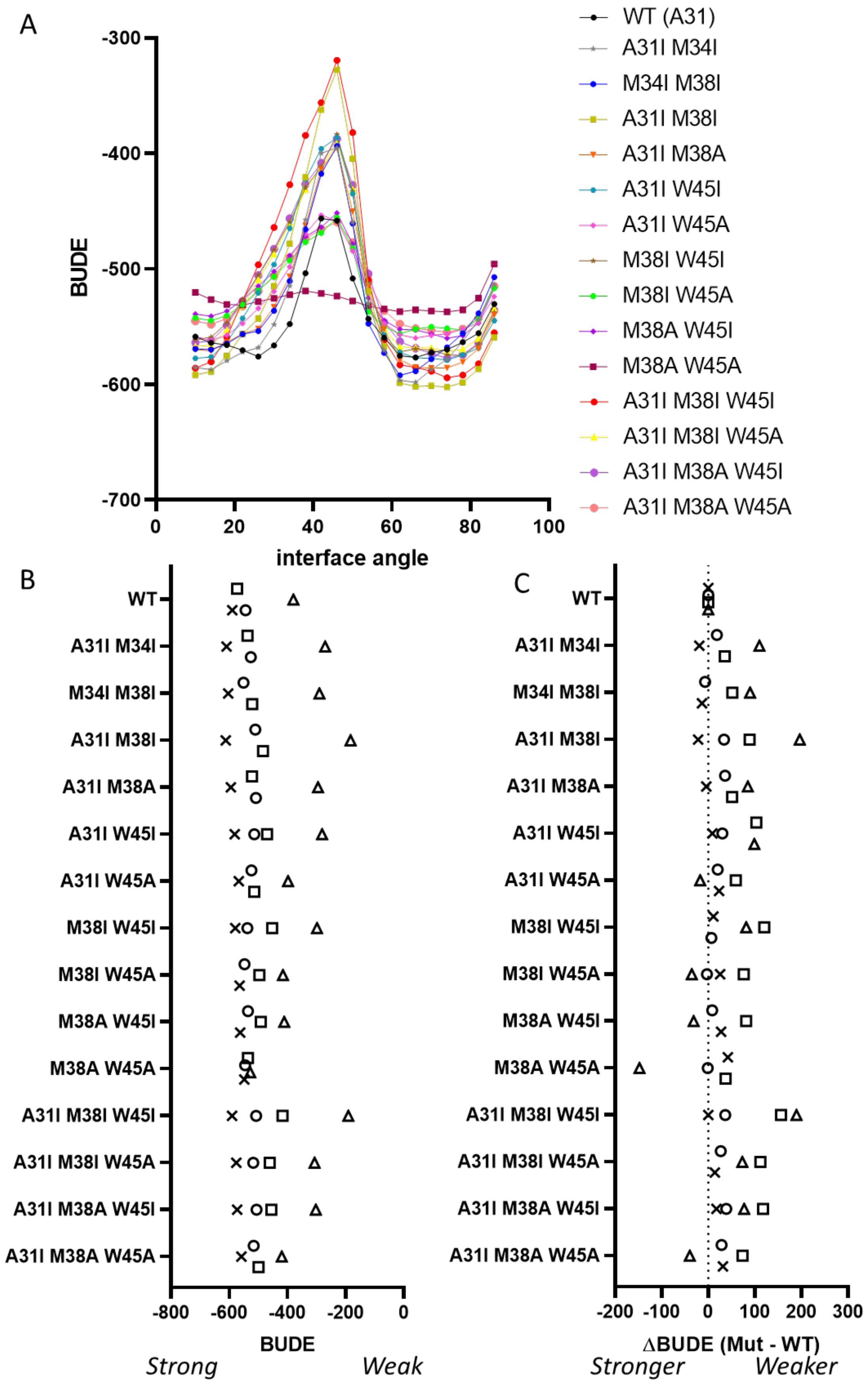

**Supplementary Figure 2. 2D BUDE profiles of all MotB variants.**

A) 2D BUDE profiles of all double- and triple-mutant variants of *Ec*MotB included in the study. WT is shown in black. Configurations at 28° (square), 44° (triangle), 56° (circle) and 64° (cross) are indicated on the profiles. B) Calculated BUDE (left) and ΔBUDE (Mutant – WT, Right) values at the 28° (square), 44° (triangle), 56° (circle) and 64° (cross) configurations for all double and triple mutants included in the study.

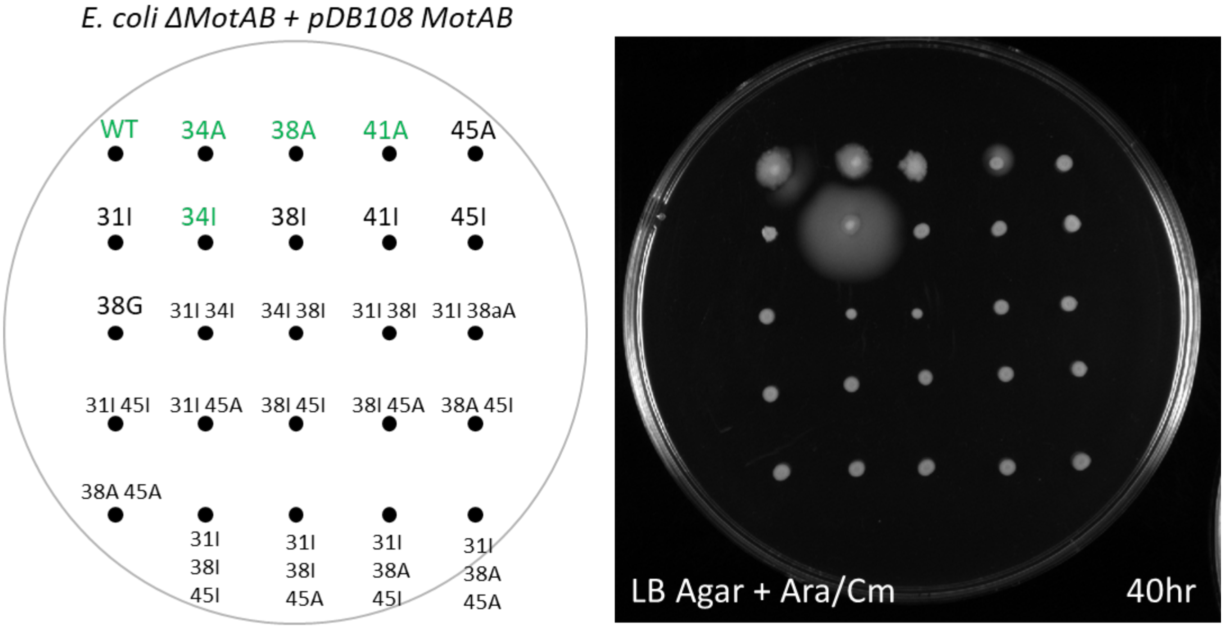

**Supplementary Figure 3. Testing MotB variants in vivo in *E. coli.***

Swim plate swim assay of WT *Ec*MotB and all variants included in the study. A pDB108 plasmid encoding the mutated variant along with WT MotA was expressed in a ΔMotAB strain *E. coli.* The schematic on the left indicates the position of each variant being tested on the swim plate on the right. The plate was incubated at 30°C for 40hr. Variants highlighted in green were motile while the ones labelled in black were not.

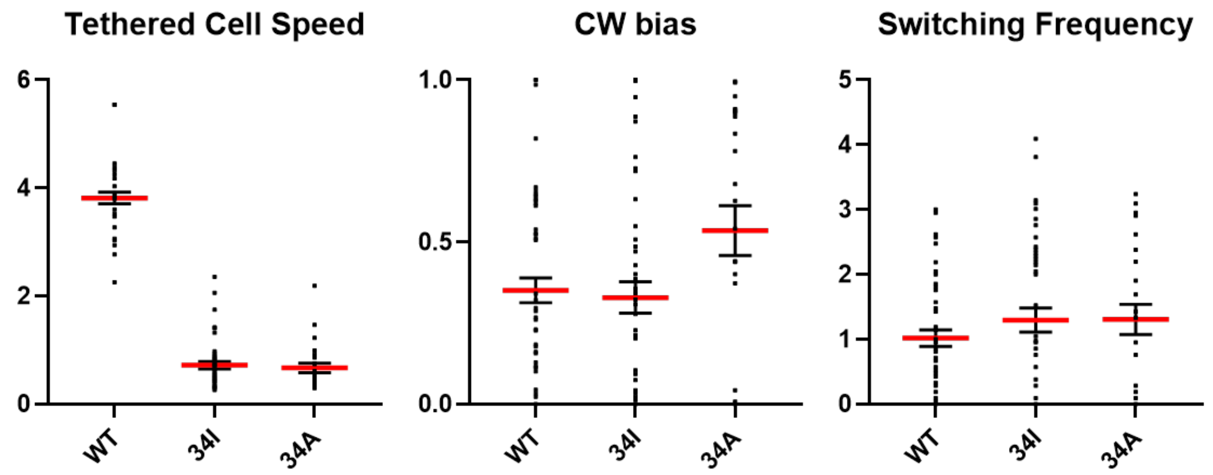

**Supplementary Figure 4. Functional characterization of WT MotB, M34I and M34A mutants.**

Tethered cell assay parameters experimentally determined for WT, M34I and M34A variants: tethered cell speed (Hz, left), ClockWise bias (CW/CCW, center) and switching frequency (1/s, right). N = 50, 46, 25 cells. Error bars indicate Mean and Standard Deviation in the Speed and Switching Frequency plots (left and right), Mean and Standard Error of the Mean (SEM) for the CW bias plot (center)

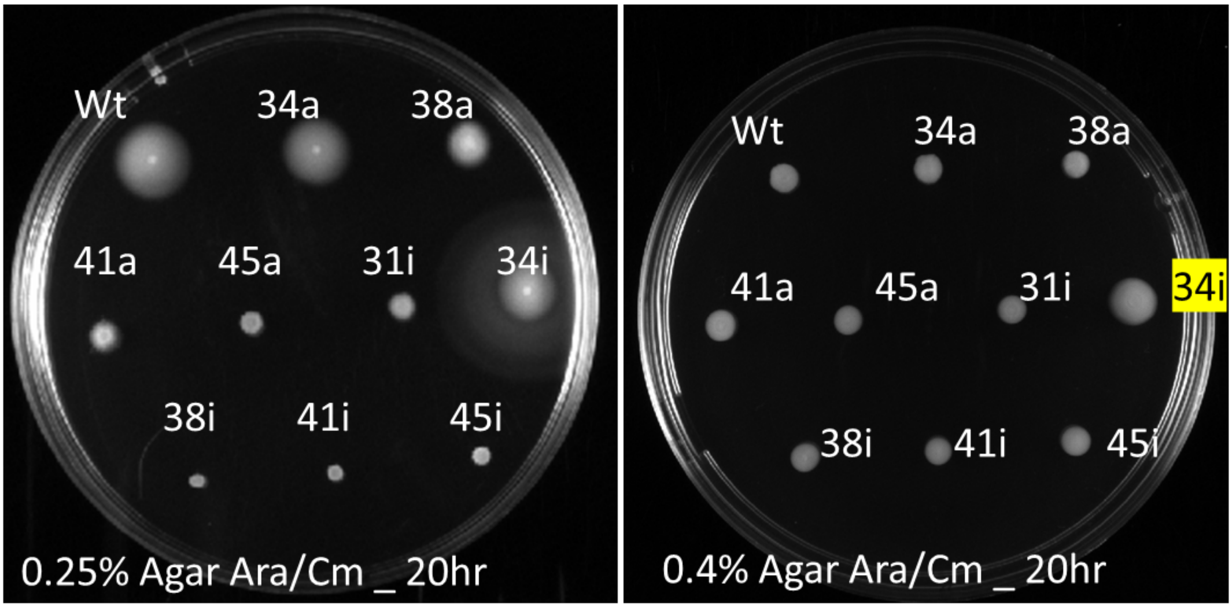

**Supplementary Figure 5. Testing MotB variants on agar swim plates of different stiffness.** Swim plate swim assay of WT *Ec*MotB and all single Alanine and Isoleucine mutants on softer (0.25% agar) or thicker agar (0.4%). Variant M34I was found to be the only mutant able to swim on the 0.4% agar swim plate and is highlighted in yellow. Plates were incubated at 30°C for 20hr.

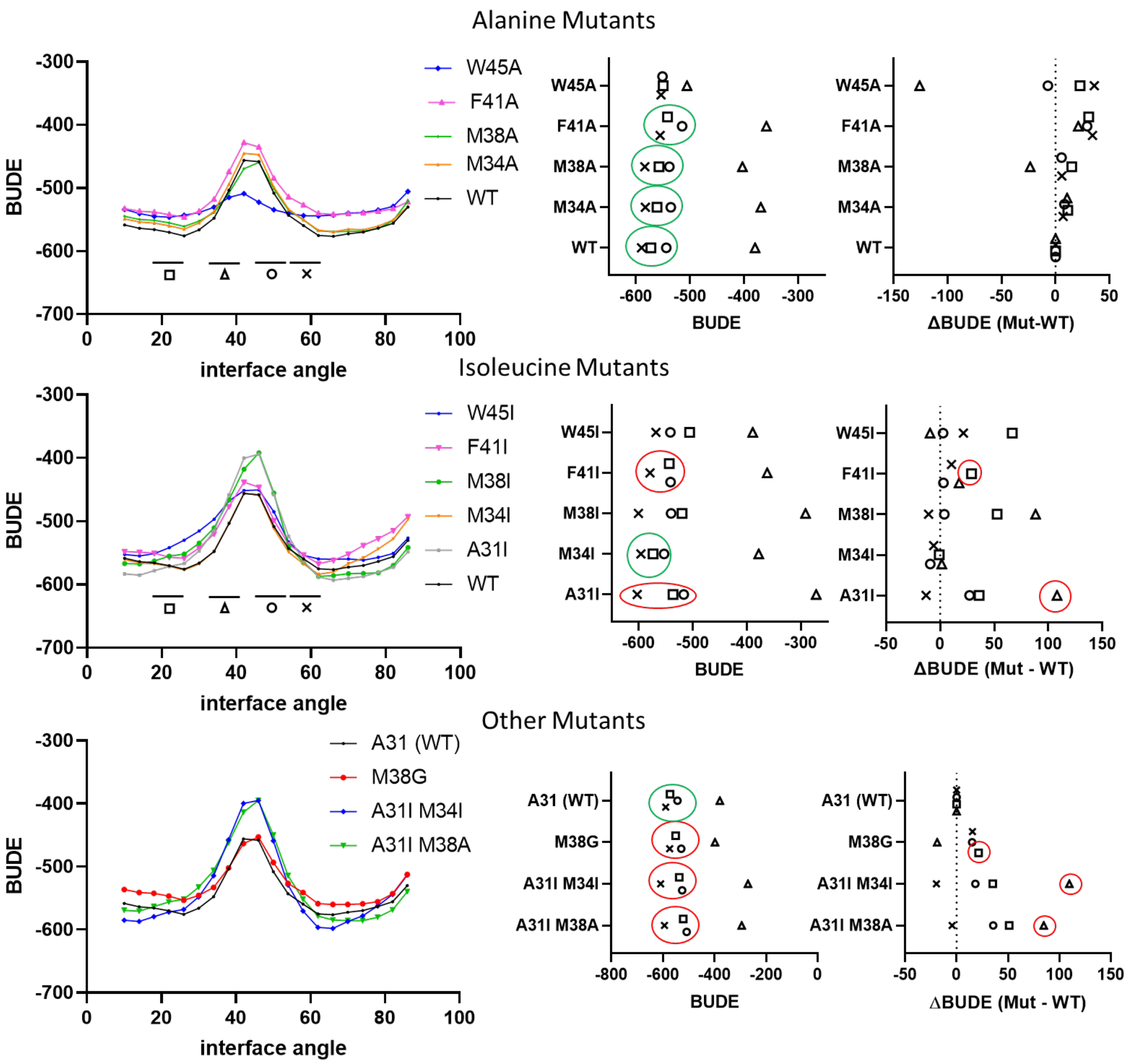

**Supplementary Figure 6. BUDE profile features correlated with motile and non-motile phenotypes.**

Two-dimensional BUDE profiles (left), BUDE values (centre) and ΔBUDE (right) compared to WT, calculated at the 28° (square), 44° (triangle), 56° (circle) and 64° (cross) configurations for Alanine (top) or Isoleucine (middle) and other (bottom) replacements at positions 31, 34, 38, 41, and 45 of WT *Ec*MotB. BUDE values graphs (centre): Green circles indicate motile variants displaying a cross-square-circle pattern of BUDE values, red circles indicate non-motile variants also displaying the same pattern. ΔBUDE graphs (right): Red circles indicate destabilized conformations (28° and 44°) in non-motile variants displaying the cross-square-circle pattern that may be causing the loss-of motility.

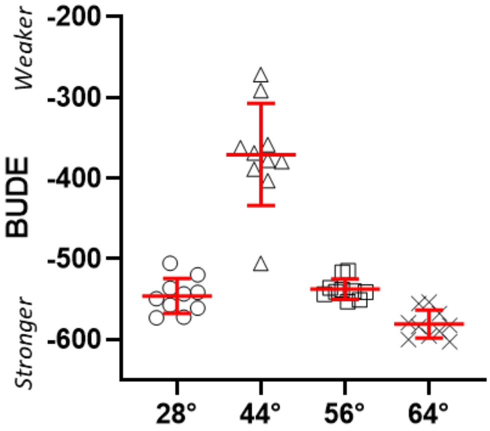

| | 28° | 56° | 64° | 44° |
|---|---|---|---|---|
| Number of values | 10 | 10 | 10 | 10 |
| Minimum | -573.1 | -553.0 | -602.4 | -505.4 |
| Maximum | -505.5 | -514.7 | -553.6 | -271.5 |
| Range | 67.60 | 38.30 | 48.80 | 233.9 |
| Mean | -545.9 | -537.5 | -580.9 | -370.7 |
| Std. Deviation | 21.73 | 12.71 | 17.32 | 63.26 |
| Std. Error of Mean | 6.872 | 4.020 | 5.477 | 20.00 |

**Supplementary Figure 7. Variability of BUDE values at each configuration hotspot.**

Average BUDE values at each hotspot configuration for WT and all single point mutants (left). Error bars indicate Mean and Standard Deviation. Complete descriptive statistics for the plotted points are provided on the right. The standard deviation of the 56° configuration is highlighted in red.

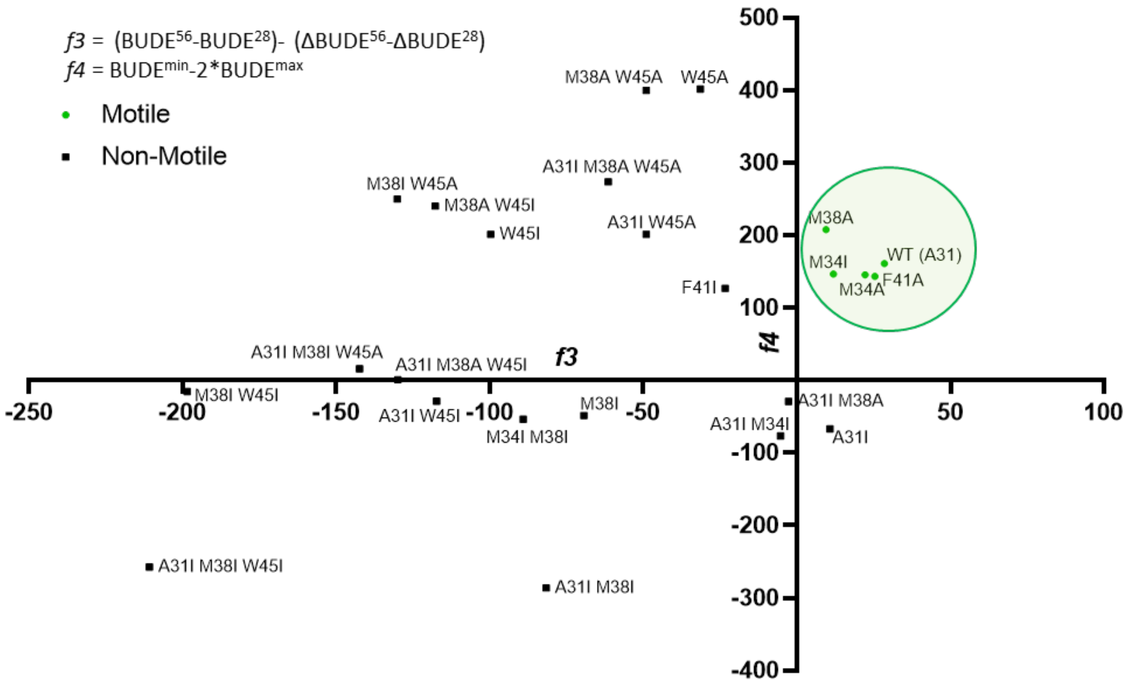

**Supplementary Figure 8. Improved parametrization of BUDE profiles for the discrimination of motile and non-motile *Ec*MotB variants**. Plot of each MotB variant mapped on an XY graph where X = *f3* and Y = *f4*. The formulas used to calculate *f3* and *f4* are provided on the top left. The green box indicates a cluster of motile variants.

A

| | |
|---|---|
| Blastochloris_sp. | VTFADLMGLLMMFFVTLTAF |
| Methylobacterium_haplocladii | VTFADLMALLMSFFVMIAAY |
| Bradyrhizobium_diazoefficiens | VTFADLMGLMMSFFVMLVAF |
| Paenibacillus | LPYSDLMTLLVALFIVMYSM |
| Moorella_thermoacetica | LTYSDLITLLMIFFVVMYAI |
| Syntrophomonas_wolfei | ITYSDLITLLMVFFVMMYSM |
| Acidobacteria bacterium1 | VAYADFVTAMMALFIVLWLM |
| Bryobacterales_bacterium | VAYADFVTAMMALFIVLWLL |
| Sodalis_glossinidius | IAYADFMTAMMAFFMVMWML |
| Vibrio_MotS | VAFADFMIALMALFLVLWVM |
| Aeromonas salmonicida1 | VAFADFTLAMMAFFMVLWIL |
| Bradyrhizobium_japonicum | IAYADFMTAMMAFFLVMWLL |
| Escherichia coli | IAYADFMTAMMAFFLVMWLI |
| Mesorhizobium | IAFADFMTAMMCFFLVMWLI |
| Hahella_chejuensis1 | VAFADFATAMMAFFLVLWLS |
| Syntrophus_aciditrophicus | VAYADFVTAMMAFFLLLWLI |
| Chromobacterium_violaceum | IAYADFVTAMMAFFLIMWLL |
| Shewanella_violacea | ATFADLMSLLMCFFILLLSF |
| Marinobacter_hydrocarbonoclasticus | VTFADLMSLLMCFFVLLLSF |
| Pseudoalteromonas_sp. | GTFADLMSLLMCFFVLLLAF |
| Hahella_chejuensis | ATFADLMSLLMCFFVLLLSF |
| Aeromonas_schubertii | GTFADLMSLLMCFFVLLLSF |
| V.Alginolyticus_PomB | GTFADLMSLLMCFFVLLLSF |
| Rhodanobacter_sp | IPYGDLLTLLLALFVVMYAV |
| Lysobacter_sp. | IPYADLLTLLLAFFVVMYAV |
| Bacillus_subtilis_subsp._subtilis_str._168 | VPYADILTLLLALFIVLYAS |
| Hydrogenobaculum_sp. | IPYADFLTLLLALFIALYAI |
| Campilobacter_jejuni | ATFSDTITLLLTFFILLYSF |
| Caldicellulosiruptor_saccharolyticus | ITYSDLITLLLIYFVVLYSM |
| Paenibacillus_odorifer | ITYADLITLLLIFFVILYAM |
| Amphibacillus_xylanus | TTYSDLVTLLLCFFVLLYSM |
| Desulfovibrio_brasiliensis | TTFADLSMLLLVFFILLYSM |
| Campilobacter_sporogenes | VPYADFLSLLLALFIALWAI |
| Marinomonas_piezotolerans | FTFADLMSLLLVFFILLYSL |
| Leptospira | LTYGDMVTLLLCFFIMLYTT |
| Lawsonia_intracellularis | TVFCDISLLLLTFFILILSM |
| Gammaproteobacteria_bacterium3 | VSYADFITLLLAFFVVMYSI |
| Aquifex_aeolicus_VF5 | TSFGDLMSLLLTFFILLYSM |
| Cellulomonas_hominis | VSYSDMITVLMALFIVLFAI |
| Marinobacter_salsuginis | VTFADLATLLLTFFILLLSF |
| Oceanobacillus_iheyensis1 | LPYADLLTLLVALFIVLFAM |
| Gammaproteobacteria_bacterium2 | VSYADFMTLLFVFFVVMYAV |
| Gammaproteobacteria_bacterium1 | ISYSDFITLLFAFFVVMYSI |
| Bdellovibrio_bacteriovorus | VSYADFITLLFAFFVVMYAT |
| Nitrospirae_bacterium | VSYADFITLLFAFFVVMYAV |
| Amphritea_balenae | VSYADFITLLFAFFVVMYAI |
| Shewanella_oneidensis | ISYADFMTLLFALFVVLYSF |
| Geobacter_sulfurreducens | VSYADFITLLFAVFVTLYAM |
| Aeromonas_salmonicida2 | VSYADFMTLLFALFVVLYSF |
| Acidobacteria_bacterium3 | VSYADFITLLFAFFVTLYAI |
| Acidithiobacillus_sulfuriphilus | VSYADFITLLFSFFVVMYAI |
| Pseudomonas | VSYADFITLLFAFFVVMYSI |
| Listeria | IPYSDLLTLLLALFIVLFAS |
| Falsibacillus_pallidus | IPYADLLTLVLALFIVLFAA |
| Heliobacterium_modesticaldum | IPYADMLTLLLALFIVMFAV |
| Bacillus_swezeyi | IPYADLLTLLLALFIVLFAM |
| Bacillus_pumilus | IPYADLLTLLLALFIVLFAS |
| Solibacillus_silvestris | VPYADILTLLLALFIVLFAS |
| Bacillus_alcalophilus | VTFSDLMTLILVFFVMLFSM |
| Bacillus_sp. | VTFSDLMTLILVFFILLFSM |
| Bacillus_pseudofirmus | VTFSDMMTLILVFFILLFSM |
| Oceanobacillus_iheyensis2 | VTYSDMVTLILVFFILLFSM |
| Bacillus_firmus | VTFSDLVTLILVFFILLFSM |
| Oceanobacillus_iheyensis3 | VTYSDMITLILVFFILLFSM |
| Bacillus_licheniformis | VTFTDLITLILVFFILLFSM |
| Treponema pallidum | TTYSDMVTLMLCFFVMLFNP |
| Bacillus_megaterium | VTFADLVTLILVFFILLFSM |
| Halodesulfovibrio_spirochaetisodalis | TTFADMVTLLLAFFVLLFSM |
| Acidobacteria_bacterium4 | VSYADFITLLFAFFVVMFAT |
| Candidatus_solibacter | VSYADFITLLFAFFVVMFAS |
| Deltaproteobacteria | VSYADFITLLFAVFVTLFAM |

B

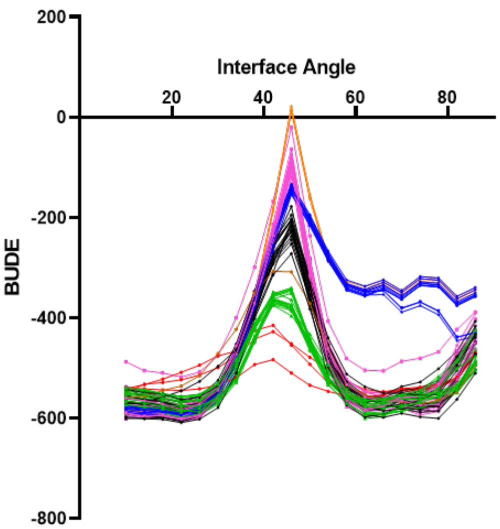

C

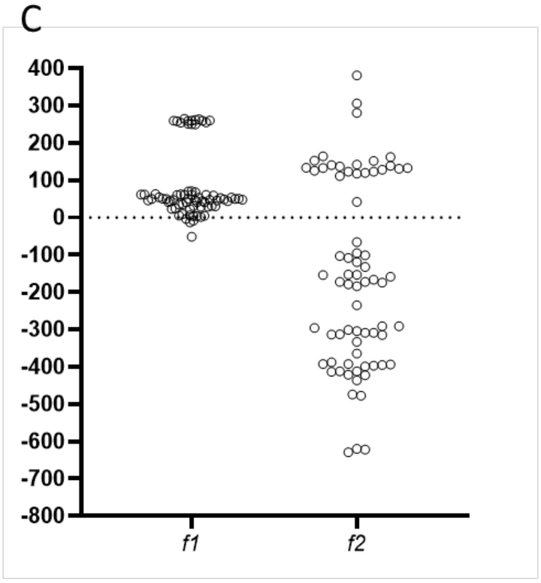

**Supplementary Figure 9. BUDE profile analysis of extant MotB homologs.**

A) List of species and respective 20 amino acid homologous sequence included in the analysis of extant MotB's, color-coded according to the clusters presented in B and Fig.3A. Notable species in the list are also highlighted by a black box.

B) 2D Bude plots for all species described in A.

C) Parameters *f1* and *f2* plotted side by side for all profiles describe in B.

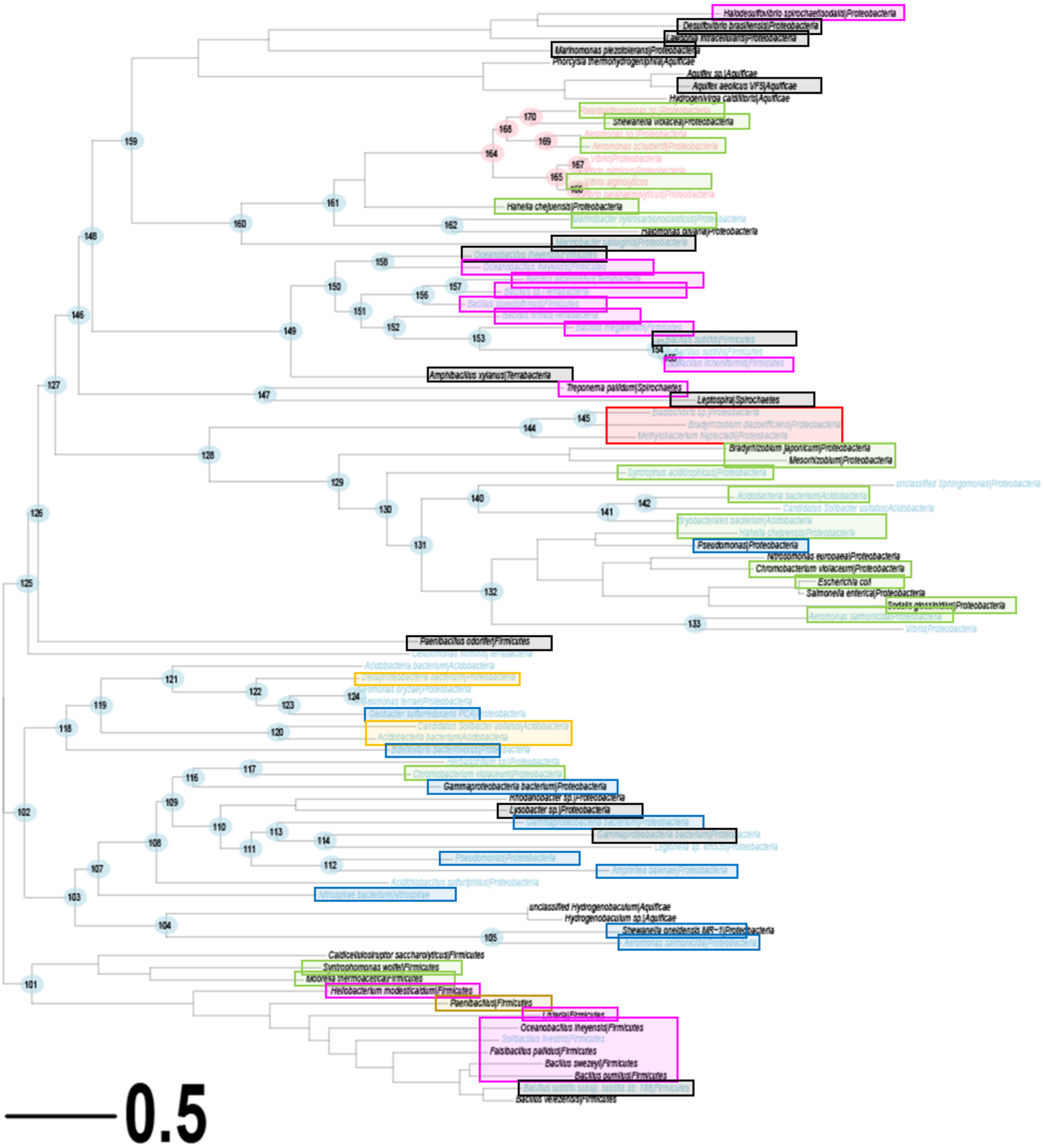

**Supplementary Figure 10. Phylogeny of MotB of 91 bacterial species.** Tips are highlighted by boxes coloured according to the clustering shown in Fig.4 and SI Fig.9.

**Supplementary Table 1 – List of MotB mutants for *in silico* modelling**

| MotB Variant | 20 residue amino acid sequence |
|---|---|
| WT *Ec*MotB | IAYADFMTAMMAFFLVMWLI |
| M34I | IAYADFITAMMAFFLVMWLI |
| M38G | IAYADFMTAMGAFFLVMWLI |
| F41I | IAYADFMTAMMAFILVMWLI |
| A31I M34I | IAYIDFITAMMAFFLVMWLI |
| M34I M38I | IAYADFITAMIAFFLVMWLI |

| | |
|---|---|
| W45I | IAYADFMTAMMAFFLVMILI |
| W45A | IAYADFMTAMMAFFLVMALI |
| F41L | IAYADFMTAMMAFLLVMWLI |
| F41L W45I | IAYADFMTAMMAFLLVMILI |
| F41L W45A | IAYADFMTAMMAFLLVMALI |
| F41A | IAYADFMTAMMAFALVMWLI |
| F41A W45I | IAYADFMTAMMAFALVMILI |
| F41A W45A | IAYADFMTAMMAFALVMALI |
| M38I | IAYADFMTAMIAFFLVMWLI |
| M38I W45I | IAYADFMTAMIAFFLVMILI |
| M38I W45A | IAYADFMTAMIAFFLVMALI |
| M38I F41L | IAYADFMTAMIAFLLVMWLI |
| M38I F41L W45I | IAYADFMTAMIAFLLVMILI |
| M38I F41L W45A | IAYADFMTAMIAFLLVMALI |
| M38I F41A | IAYADFMTAMIAFALVMWLI |
| M38I F41A W45I | IAYADFMTAMIAFALVMILI |
| M38I F41A W45A | IAYADFMTAMIAFALVMALI |
| M38N | IAYADFMTAMNAFFLVMWLI |
| M38N W45I | IAYADFMTAMNAFFLVMILI |
| M38N W45A | IAYADFMTAMNAFFLVMALI |
| M38N F41L | IAYADFMTAMNAFLLVMWLI |
| M38N F41L W45I | IAYADFMTAMNAFLLVMILI |
| M38N F41L W45A | IAYADFMTAMNAFLLVMALI |
| M38N F41A | IAYADFMTAMNAFALVMWLI |
| M38N F41A W45I | IAYADFMTAMNAFALVMILI |
| M38N F41A W45A | IAYADFMTAMNAFALVMALI |
| M38A | IAYADFMTAMAAFFLVMWLI |
| M38A W45I | IAYADFMTAMAAFFLVMILI |
| M38A W45A | IAYADFMTAMAAFFLVMALI |
| M38A F41L | IAYADFMTAMAAFLLVMWLI |
| M38A F41L W45I | IAYADFMTAMAAFLLVMILI |
| M38A F41L W45A | IAYADFMTAMAAFLLVMALI |
| M38A F41A | IAYADFMTAMAAFALVMWLI |
| M38A F41A W45I | IAYADFMTAMAAFALVMILI |
| M38A F41A W45A | IAYADFMTAMAAFALVMALI |
| M34L | IAYADFLTAMMAFFLVMWLI |
| M34L W45I | IAYADFLTAMMAFFLVMILI |
| M34L W45A | IAYADFLTAMMAFFLVMALI |
| M34L F41L | IAYADFLTAMMAFLLVMWLI |
| M34L F41L W45I | IAYADFLTAMMAFLLVMILI |
| M34L F41L W45A | IAYADFLTAMMAFLLVMALI |
| M34L F41A | IAYADFLTAMMAFALVMWLI |
| M34L F41A W45I | IAYADFLTAMMAFALVMILI |
| M34L F41A W45A | IAYADFLTAMMAFALVMALI |
| M34L M38I | IAYADFLTAMIAFFLVMWLI |
| M34L M38I W45I | IAYADFLTAMIAFFLVMILI |

| | |
|---|---|
| M34L M38I W45A | IAYADFLTAMIAFFLVMALI |
| M34L M38I F41L | IAYADFLTAMIAFLLVMWLI |
| M34L M38I F41L W45I | IAYADFLTAMIAFLLVMILI |
| M34L M38I F41L W45A | IAYADFLTAMIAFLLVMALI |
| M34L M38I F41A | IAYADFLTAMIAFALVMWLI |
| M34L M38I F41A W45I | IAYADFLTAMIAFALVMILI |
| M34L M38I F41A W45A | IAYADFLTAMIAFALVMALI |
| M34L M38N | IAYADFLTAMNAFFLVMWLI |
| M34L M38N W45I | IAYADFLTAMNAFFLVMILI |
| M34L M38N W45A | IAYADFLTAMNAFFLVMALI |
| M34L M38N F41L | IAYADFLTAMNAFLLVMWLI |
| M34L M38N F41L W45I | IAYADFLTAMNAFLLVMILI |
| M34L M38N F41L W45A | IAYADFLTAMNAFLLVMALI |
| M34L M38N F41A | IAYADFLTAMNAFALVMWLI |
| M34L M38N F41A W45I | IAYADFLTAMNAFALVMILI |
| M34L M38N F41A W45A | IAYADFLTAMNAFALVMALI |
| M34L M38A | IAYADFLTAMAAFFLVMWLI |
| M34L M38A W45I | IAYADFLTAMAAFFLVMILI |
| M34L M38A W45A | IAYADFLTAMAAFFLVMALI |
| M34L M38A F41L | IAYADFLTAMAAFLLVMWLI |
| M34L M38A F41L W45I | IAYADFLTAMAAFLLVMILI |
| M34L M38A F41L W45A | IAYADFLTAMAAFLLVMALI |
| M34L M38A F41A | IAYADFLTAMAAFALVMWLI |
| M34L M38A F41A W45I | IAYADFLTAMAAFALVMILI |
| M34L M38A F41A W45A | IAYADFLTAMAAFALVMALI |
| M34A | IAYADFATAMMAFFLVMWLI |
| M34A W45I | IAYADFATAMMAFFLVMILI |
| M34A W45A | IAYADFATAMMAFFLVMALI |
| M34A F41L | IAYADFATAMMAFLLVMWLI |
| M34A F41L W45I | IAYADFATAMMAFLLVMILI |
| M34A F41L W45A | IAYADFATAMMAFLLVMALI |
| M34A F41A | IAYADFATAMMAFALVMWLI |
| M34A F41A W45I | IAYADFATAMMAFALVMILI |
| M34A F41A W45A | IAYADFATAMMAFALVMALI |
| M34A M38I | IAYADFATAMIAFFLVMWLI |
| M34A M38I W45I | IAYADFATAMIAFFLVMILI |
| M34A M38I W45A | IAYADFATAMIAFFLVMALI |
| M34A M38I F41L | IAYADFATAMIAFLLVMWLI |
| M34A M38I F41L W45I | IAYADFATAMIAFLLVMILI |
| M34A M38I F41L W45A | IAYADFATAMIAFLLVMALI |
| M34A M38I F41A | IAYADFATAMIAFALVMWLI |
| M34A M38I F41A W45I | IAYADFATAMIAFALVMILI |
| M34A M38I F41A W45A | IAYADFATAMIAFALVMALI |
| M34A M38N | IAYADFATAMNAFFLVMWLI |
| M34A M38N W45I | IAYADFATAMNAFFLVMILI |
| M34A M38N W45A | IAYADFATAMNAFFLVMALI |

| | |
|---|---|
| M34A M38N F41L | IAYADFATAMNAFLLVMWLI |
| M34A M38N F41L W45I | IAYADFATAMNAFLLVMILI |
| M34A M38N F41L W45A | IAYADFATAMNAFLLVMALI |
| M34A M38N F41A | IAYADFATAMNAFALVMWLI |
| M34A M38N F41A W45I | IAYADFATAMNAFALVMILI |
| M34A M38N F41A W45A | IAYADFATAMNAFALVMALI |
| M34A M38A | IAYADFATAMAAFFLVMWLI |
| M34A M38A W45I | IAYADFATAMAAFFLVMILI |
| M34A M38A W45A | IAYADFATAMAAFFLVMALI |
| M34A M38A F41L | IAYADFATAMAAFLLVMWLI |
| M34A M38A F41L W45I | IAYADFATAMAAFLLVMILI |
| M34A M38A F41L W45A | IAYADFATAMAAFLLVMALI |
| M34A M38A F41A | IAYADFATAMAAFALVMWLI |
| M34A M38A F41A W45I | IAYADFATAMAAFALVMILI |
| M34A M38A F41A W45A | IAYADFATAMAAFALVMALI |
| A31I | IAYIDFMTAMMAFFLVMWLI |
| A31I W45I | IAYIDFMTAMMAFFLVMILI |
| A31I W45A | IAYIDFMTAMMAFFLVMALI |
| A31I F41L | IAYIDFMTAMMAFLLVMWLI |
| A31I F41L W45I | IAYIDFMTAMMAFLLVMILI |
| A31I F41L W45A | IAYIDFMTAMMAFLLVMALI |
| A31I F41A | IAYIDFMTAMMAFALVMWLI |
| A31I F41A W45I | IAYIDFMTAMMAFALVMILI |
| A31I F41A W45A | IAYIDFMTAMMAFALVMALI |
| A31I M38I | IAYIDFMTAMIAFFLVMWLI |
| A31I M38I W45I | IAYIDFMTAMIAFFLVMILI |
| A31I M38I W45A | IAYIDFMTAMIAFFLVMALI |
| A31I M38I F41L | IAYIDFMTAMIAFLLVMWLI |
| A31I M38I F41L W45I | IAYIDFMTAMIAFLLVMILI |
| A31I M38I F41L W45A | IAYIDFMTAMIAFLLVMALI |
| A31I M38I F41A | IAYIDFMTAMIAFALVMWLI |
| A31I M38I F41A W45I | IAYIDFMTAMIAFALVMILI |
| A31I M38I F41A W45A | IAYIDFMTAMIAFALVMALI |
| A31I M38N | IAYIDFMTAMNAFFLVMWLI |
| A31I M38N W45I | IAYIDFMTAMNAFFLVMILI |
| A31I M38N W45A | IAYIDFMTAMNAFFLVMALI |
| A31I M38N F41L | IAYIDFMTAMNAFLLVMWLI |
| A31I M38N F41L W45I | IAYIDFMTAMNAFLLVMILI |
| A31I M38N F41L W45A | IAYIDFMTAMNAFLLVMALI |
| A31I M38N F41A | IAYIDFMTAMNAFALVMWLI |
| A31I M38N F41A W45I | IAYIDFMTAMNAFALVMILI |
| A31I M38N F41A W45A | IAYIDFMTAMNAFALVMALI |
| A31I M38A | IAYIDFMTAMAAFFLVMWLI |
| A31I M38A W45I | IAYIDFMTAMAAFFLVMILI |
| A31I M38A W45A | IAYIDFMTAMAAFFLVMALI |
| A31I M38A F41L | IAYIDFMTAMAAFLLVMWLI |

| | |
|---|---|
| A31I M38A F41L W45I | IAYIDFMTAMAAFLLVMILI |
| A31I M38A F41L W45A | IAYIDFMTAMAAFLLVMALI |
| A31I M38A F41A | IAYIDFMTAMAAFALVMWLI |
| A31I M38A F41A W45I | IAYIDFMTAMAAFALVMILI |
| A31I M38A F41A W45A | IAYIDFMTAMAAFALVMALI |
| A31I M34L | IAYIDFLTAMMAFFLVMWLI |
| A31I M34L W45I | IAYIDFLTAMMAFFLVMILI |
| A31I M34L W45A | IAYIDFLTAMMAFFLVMALI |
| A31I M34L F41L | IAYIDFLTAMMAFLLVMWLI |
| A31I M34L F41L W45I | IAYIDFLTAMMAFLLVMILI |
| A31I M34L F41L W45A | IAYIDFLTAMMAFLLVMALI |
| A31I M34L F41A | IAYIDFLTAMMAFALVMWLI |
| A31I M34L F41A W45I | IAYIDFLTAMMAFALVMILI |
| A31I M34L F41A W45A | IAYIDFLTAMMAFALVMALI |
| A31I M34L M38I | IAYIDFLTAMIAFFLVMWLI |
| A31I M34L M38I W45I | IAYIDFLTAMIAFFLVMILI |
| A31I M34L M38I W45A | IAYIDFLTAMIAFFLVMALI |
| A31I M34L M38I F41L | IAYIDFLTAMIAFLLVMWLI |
| A31I M34L M38I F41L W45I | IAYIDFLTAMIAFLLVMILI |
| A31I M34L M38I F41L W45A | IAYIDFLTAMIAFLLVMALI |
| A31I M34L M38I F41A | IAYIDFLTAMIAFALVMWLI |
| A31I M34L M38I F41A W45I | IAYIDFLTAMIAFALVMILI |
| A31I M34L M38I F41A W45A | IAYIDFLTAMIAFALVMALI |
| A31I M34L M38N | IAYIDFLTAMNAFFLVMWLI |
| A31I M34L M38N W45I | IAYIDFLTAMNAFFLVMILI |
| A31I M34L M38N W45A | IAYIDFLTAMNAFFLVMALI |
| A31I M34L M38N F41L | IAYIDFLTAMNAFLLVMWLI |
| A31I M34L M38N F41L W45I | IAYIDFLTAMNAFLLVMILI |
| A31I M34L M38N F41L W45A | IAYIDFLTAMNAFLLVMALI |
| A31I M34L M38N F41A | IAYIDFLTAMNAFALVMWLI |
| A31I M34L M38N F41A W45I | IAYIDFLTAMNAFALVMILI |
| A31I M34L M38N F41A W45A | IAYIDFLTAMNAFALVMALI |
| A31I M34L M38A | IAYIDFLTAMAAFFLVMWLI |
| A31I M34L M38A W45I | IAYIDFLTAMAAFFLVMILI |
| A31I M34L M38A W45A | IAYIDFLTAMAAFFLVMALI |
| A31I M34L M38A F41L | IAYIDFLTAMAAFLLVMWLI |
| A31I M34L M38A F41L W45I | IAYIDFLTAMAAFLLVMILI |
| A31I M34L M38A F41L W45A | IAYIDFLTAMAAFLLVMALI |
| A31I M34L M38A F41A | IAYIDFLTAMAAFALVMWLI |
| A31I M34L M38A F41A W45I | IAYIDFLTAMAAFALVMILI |
| A31I M34L M38A F41A W45A | IAYIDFLTAMAAFALVMALI |
| A31I M34A | IAYIDFATAMMAFFLVMWLI |
| A31I M34A W45I | IAYIDFATAMMAFFLVMILI |
| A31I M34A W45A | IAYIDFATAMMAFFLVMALI |
| A31I M34A F41L | IAYIDFATAMMAFLLVMWLI |
| A31I M34A F41L W45I | IAYIDFATAMMAFLLVMILI |

| | |
|---|---|
| A31I M34A F41L W45A | IAYIDFATAMMAFLLVMALI |
| A31I M34A F41A | IAYIDFATAMMAFALVMWLI |
| A31I M34A F41A W45I | IAYIDFATAMMAFALVMILI |
| A31I M34A F41A W45A | IAYIDFATAMMAFALVMALI |
| A31I M34A M38I | IAYIDFATAMIAFFLVMWLI |
| A31I M34A M38I W45I | IAYIDFATAMIAFFLVMILI |
| A31I M34A M38I W45A | IAYIDFATAMIAFFLVMALI |
| A31I M34A M38I F41L | IAYIDFATAMIAFLLVMWLI |
| A31I M34A M38I F41L W45I | IAYIDFATAMIAFLLVMILI |
| A31I M34A M38I F41L W45A | IAYIDFATAMIAFLLVMALI |
| A31I M34A M38I F41A | IAYIDFATAMIAFALVMWLI |
| A31I M34A M38I F41A W45I | IAYIDFATAMIAFALVMILI |
| A31I M34A M38I F41A W45A | IAYIDFATAMIAFALVMALI |
| A31I M34A M38N | IAYIDFATAMNAFFLVMWLI |
| A31I M34A M38N W45I | IAYIDFATAMNAFFLVMILI |
| A31I M34A M38N W45A | IAYIDFATAMNAFFLVMALI |
| A31I M34A M38N F41L | IAYIDFATAMNAFLLVMWLI |
| A31I M34A M38N F41L W45I | IAYIDFATAMNAFLLVMILI |
| A31I M34A M38N F41L W45A | IAYIDFATAMNAFLLVMALI |
| A31I M34A M38N F41A | IAYIDFATAMNAFALVMWLI |
| A31I M34A M38N F41A W45I | IAYIDFATAMNAFALVMILI |
| A31I M34A M38N F41A W45A | IAYIDFATAMNAFALVMALI |
| A31I M34A M38A | IAYIDFATAMAAFFLVMWLI |
| A31I M34A M38A W45I | IAYIDFATAMAAFFLVMILI |
| A31I M34A M38A W45A | IAYIDFATAMAAFFLVMALI |
| A31I M34A M38A F41L | IAYIDFATAMAAFLLVMWLI |
| A31I M34A M38A F41L W45I | IAYIDFATAMAAFLLVMILI |
| A31I M34A M38A F41L W45A | IAYIDFATAMAAFLLVMALI |
| A31I M34A M38A F41A | IAYIDFATAMAAFALVMWLI |
| A31I M34A M38A F41A W45I | IAYIDFATAMAAFALVMILI |
| A31I M34A M38A F41A W45A | IAYIDFATAMAAFALVMALI |

**Supplementary Table 2 – List of primers for MotB mutagenesis**

| Primer name | Sequence (5' to 3') |
|---|---|
| A31I -Fw | gcacatggatcgtggaagattgcttatatcgactttatgactgcg |
| A31I-Rv | cgcagtcataaagtcgatataagcaatcttccacgatccatgtgc |
| M34A-Fw | agattgcttatgccgactttgcgactgcgatgatggccttt |
| M34A-Rv | aaaggccatcatcgcagtcgcaaagtcggcataagcaatct |
| M34I-Fw | ggaagattgcttatgccgactttataactgcgatgatgg |
| M34I-Rv | ccatcatcgcagttataaagtcggcataagcaatcttcc |

| | |
|---|---|
| M38I-Fw | ccgactttatgactgcgatgatagcctttttttctgg |
| M38I-Rv | ccagaaaaaaaggctatcatcgcagtcataaagtcgg |
| M38A-Fw | cgactttatgactgcgatggcggcctttttttctggtgatg |
| M38A-Rv | catcaccagaaaaaaaggccgccatcgcagtcataaagtcg |
| M38G-Fw | ccgactttatgactgcgatgatagcctttttttctgg |
| M38G-Rv | ccagaaaaaaaggctatcatcgcagtcataaagtcgg |
| F41A-Fw | ttatgactgcgatgatggcctttgctctggtgatgtggct |
| F41A-Rv | agccacatcaccagagcaaaggccatcatcgcagtcataa |
| F41I-Fw | ctgcgatgatggcctttattctggtgatgtggct |
| F41I-Rv | agccacatcaccagaataaaggccatcatcgcag |
| W45A-Fw | gcctttttttctggtgatggcgctgatctccatctccag |
| W45A-Rv | ctggagatggagatcagcgccatcaccagaaaaaaaggc |
| W45I-Fw | tggcctttttttctggtgatgatactgatctccatctccagccc |
| W45I-Rv | gggctggagatggagatcagtatcatcaccagaaaaaaaggcca |
| A31I/M34I-Fw | gtggaagattgcttatatcgactttataactgcgatgatggc |
| A31I/M34I-Rv | gccatcatcgcagttataaagtcgatataagcaatcttccac |
| M34I/M38I-Fw | ggaagattgcttatgccgactttataactgcgatgatag |
| M34I/M38I-Rv | ctatcatcgcagttataaagtcggcataagcaatcttcc |
